# Dissecting the roles of GRK2 and GRK3 in μ-opioid receptor internalization and β-arrestin2 recruitment using CRISPR/Cas9-edited HEK293 cells

**DOI:** 10.1101/2020.01.08.898338

**Authors:** Thor C. Møller, Mie F. Pedersen, Jeffrey R. van Senten, Sofie D. Seiersen, Jesper M. Mathiesen, Michel Bouvier, Hans Bräuner-Osborne

## Abstract

Most G protein-coupled receptors (GPCRs) recruit β-arrestins and internalize upon agonist stimulation. For the μ-opioid receptor (μ-OR), this process has been linked to development of opioid tolerance. GPCR kinases (GRKs), particularly GRK2 and GRK3, have been shown to be important for μ-OR recruitment of β-arrestin and internalization. However, the contribution of GRK2 and GRK3 to β-arrestin recruitment and receptor internalization, remain to be determined in their complete absence. Using CRISPR/Cas9-mediated genome editing we established HEK293 cells with knockout of GRK2, GRK3 or both to dissect their individual contributions in β-arrestin2 recruitment and μ-OR internalization upon stimulation with four different agonists. We showed that GRK2/3 removal reduced agonist-induced μ-OR internalization and β-arrestin2 recruitment substantially and we found GRK2 to be more important for these processes than GRK3. Furthermore, we observed a sustained and GRK2/3 independent component of β-arrestin2 recruitment to the plasma membrane upon μ-OR activation. Rescue expression experiments restored GRK2/3 functions. Inhibition of GRK2/3 using the small molecule inhibitor CMPD101 showed a high similarity between the genetic and pharmacological approaches, cross-validating the specificity of both. However, off-target effects were observed at high CMPD101 concentrations. These GRK2/3 KO cell lines should prove useful for a wide range of studies on GPCR function.

## Introduction

The family of G protein-coupled receptors (GPCRs) constitute important drug targets, through which ~30% of all clinically approved medicines mediate their action^1^. Regulation of GPCR signaling following receptor activation is a complex process that typically involves recruitment of kinases that phosphorylate the receptor to increase the receptor affinity for β-arrestins and mediate receptor internalization^2^. The μ-opioid receptor (μ-OR) belongs to the family of rhodopsin-like (family A) GPCRs and mediates the analgesic effects of opioid drugs, but also related side-effects and addictive properties^3,4^. The regulation of μ-OR desensitization and trafficking has been suggested to be linked to the development of tolerance after chronic use of opioid drugs^5^ and it is therefore important to understand the underlying molecular mechanisms.

Several studies indicate that receptor phosphorylation by GPCR kinases (GRKs) is important for the desensitization of μ-OR signaling and initiation of internalization^6–12^. Out of the seven isotypes in the GRK family, four (GRK2, GRK3, GRK5 and GRK6) have been speculated to regulate μ-OR *in vivo* due to their overlapping expression patterns^13^. Knockout (KO) models have confirmed the importance of the individual GRKs. For instance, it has been demonstrated that fentanyl and morphine-induced tolerance are decreased in GRK3 KO mice^14^, morphine reward and dependence are lost in mice depleted of GRK5, but not GRK3, and morphine-induced locomotor activity is increased in mice lacking GRK6 compared to wild type littermates^15,16^. Altogether, these studies suggest that phosphorylation of μ-OR by specific GRK subtypes differentially impacts the physiological outcome upon stimulation with opioids. The role of GRK2 has not been addressed in KO systems due to lethal effects of removing GRK2 in mouse embryos^17^. Instead, the role of GRK2 has been studied using e.g. perfusion of a GRK2-inhibitory peptide or overexpression of a GRK2 dominant negative mutant in rat neurons, both demonstrating a role of GRK2 in μ-OR desensitization^18,19^.

Further insights into the mechanisms behind the role of GRKs in μ-OR pharmacology have been obtained from *in vitro* studies. An early study demonstrated that phosphorylation of the μ- OR could be increased by GRK2 overexpression, which led to increased β-arrestin recruitment and μ- OR internalization^20^. The involvement of μ-OR phosphorylation in these processes was confirmed by later studies^10,21–24^. In addition to GRK2, *in vitro* studies have shown that GRK3/5/6 have direct roles in μ-OR phosphorylation and/or internalization^16,23,25–27^. Moreover, studies have also utilized phospho-site specific antibodies to demonstrate that the μ-OR is differentially phosphorylated by distinct GRK isotypes depending on the agonist used. For instance, stimulation with the full agonist D-Ala(2)-mephe(4)-gly-ol(5))enkephalin (DAMGO) leads to phosphorylation of T370, S375, T376 and T379 in mouse μ-OR whereas stimulation with morphine – a full agonist for G protein signaling, but a partial agonist for β-arrestin recruitment and μ-OR internalization^25^ – only leads to phosphorylation of S375^23,26^. In the same studies, the relative contribution of GRK2/3/5/6 to μ-OR phosphorylation was also dependent on the agonist used.

The tools to study the involvement of GRKs in cell systems, have for now been restricted mainly to short-interfering RNA techniques^10,23,26^, the usage of dominant negative mutants of GRK2^10,20,28^ and utilization of Takeda compound 101 (CMPD101), a reported GRK2/3 selective inhibitor^10,29–31^. To the best of our knowledge, no studies have investigated the role of GRKs in μ- OR internalization in cell systems with complete KO of specific GRKs, which would prevent incomplete knockdown of expression, residual kinase activity or unwanted overexpression effects.

Here, we report construction of individual and double KO of GRK2 and GRK3 cell lines in the human embryonic kidney 293A (HEK293A) background, which we employ to investigate the agonist-induced μ-OR internalization and β-arrestin2 recruitment in response to four different agonists. We find that KO of GRK2 and GRK3 reduces agonist-induced μ-OR internalization and β-arrestin2 recruitment. Additionally, we used a combination of β-arrestin2 recruitment assays that either report β-arrestin2 proximity to μ-OR or to the cell membrane to identify a sustained and GRK2/3 independent β-arrestin2 recruitment to the cell membrane. Furthermore, we present a side-by-side comparison of the effects obtained with CMPD101 to the responses obtained in GRK2/3 double KO cells. We find highly similar results with the two approaches when 10 μM of CMPD101 is used; higher CMPD101 concentrations lead to non-GRK2/3-mediated effects. The cell lines provide full KO of two important regulators of GPCR function and we expect them to be useful tools for future studies of GPCR function.

## Results

### Validation of GRK2 and/or GRK3 genome-edited HEK293A cells

Construction of individual and combined GRK2 and GRK3 KO HEK293A cells (ΔGRK2, ΔGRK3 and ΔGRK2/3) were performed using the clustered regularly interspaced short palindromic repeats (CRISPR)/CRISPR associated protein 9 (Cas9) technology. Clones containing modification of all alleles were identified with insertions or deletions in the *ADRBK1* locus (GRK2) and/or the *ADRBK2* locus (GRK3) leading to frameshifts as shown by sequencing and indel detection by amplicon analysis (IDAA)^32^. Furthermore, we confirmed the absence of full-length GRK2 protein expression in the ΔGRK2 and ΔGRK2/3 cells and of full-length GRK3 protein expression in the ΔGRK3 and ΔGRK2/3 cells using GRK2- and GRK3-selective antibodies (**Fig. 1**). No alteration of GRK2 expression in the ΔGRK3 clone or GRK3 expression in the ΔGRK2 clone compared to parental cells could be detected (**Fig. 1**). Similarly, the expression of GRK5 and GRK6 was comparable in the parental cells and the three ΔGRK cell lines (**Supplementary Fig. S2**).

**Figure 1.**
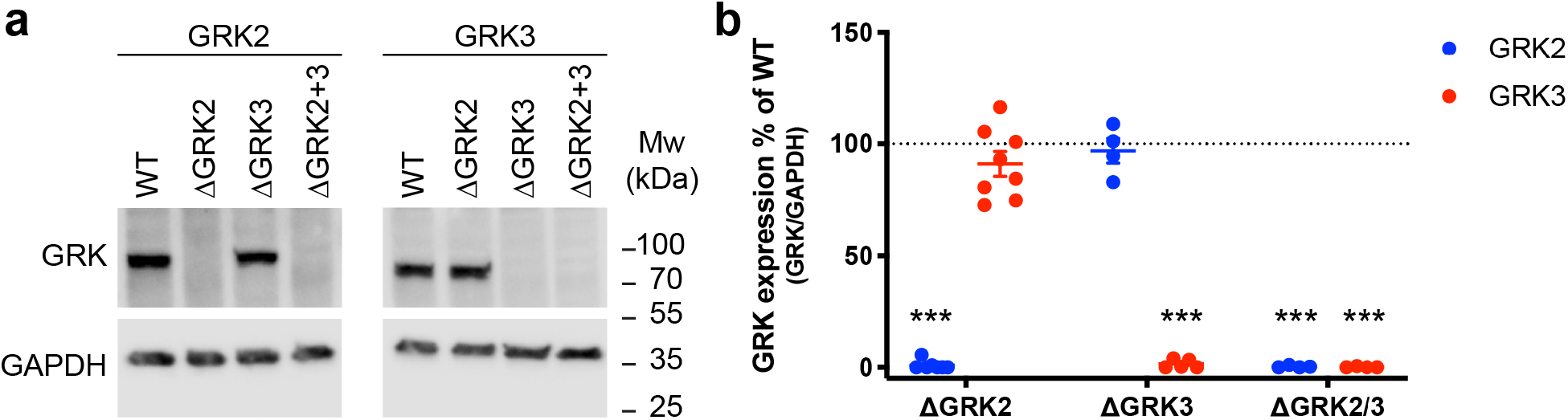
Validation of CRISPR/Cas9 knock-out cell lines. (**a**) Western blot analysis of full-length GRK2 or GRK3 expression in parental or genome-edited cell lines with deletion of GRK2, GRK3 or GRK2/3. The anti-GRK2 and anti-GRK3 antibodies target a site C-terminal to the site that is genetically modified. GAPDH expression is detected to ensure equal loading. Full-length blots are presented in Supplementary Figure 1. (**b**) Quantification of anti-GRK2 and anti-GRK3 western blots after normalization to the GAPDH signal. Mean ± SEM of 4-8 independent experiments. Expression in each of the KO cell lines was compared to the parental cells by one sample t-test using a reference value of 100 and the cutoff for what was considered a significant difference was adjusted to *P* < 0.05/n (n corresponds to the number of observations) to correct for multiple comparisons. \*\*\**P* < 0.001/n.

### GRK2 and GRK3 contributes to μ-OR internalization

We used a real-time internalization assay to determine the influence of GRK2 and GRK3 on μ-OR internalization. This assay is based on time-resolved Förster resonance energy transfer (TR-FRET) between a long lifetime donor (Lumi4-Tb) covalently linked to an amino-terminal SNAP-tag on cell surface μ-OR and a cell impermeant acceptor (fluorescein) in the extracellular buffer^33,34^.

Internalization separates the donor and acceptor molecules thus preventing energy transfer and increasing the ratio of donor over acceptor emissions (internalization ratio). We compared the ability of four μ-OR agonists to stimulate internalization: DAMGO, loperamide, fentanyl and morphine. All four agonists induced internalization in a concentration-dependent manner and reached a plateau between 45 and 90 min after agonist addition in the parental HEK293A cells as well as in the ΔGRK2, ΔGRK3 and ΔGRK2/3 cell lines (**Fig. 2, Supplementary Fig. S3**).

**Figure 2.**
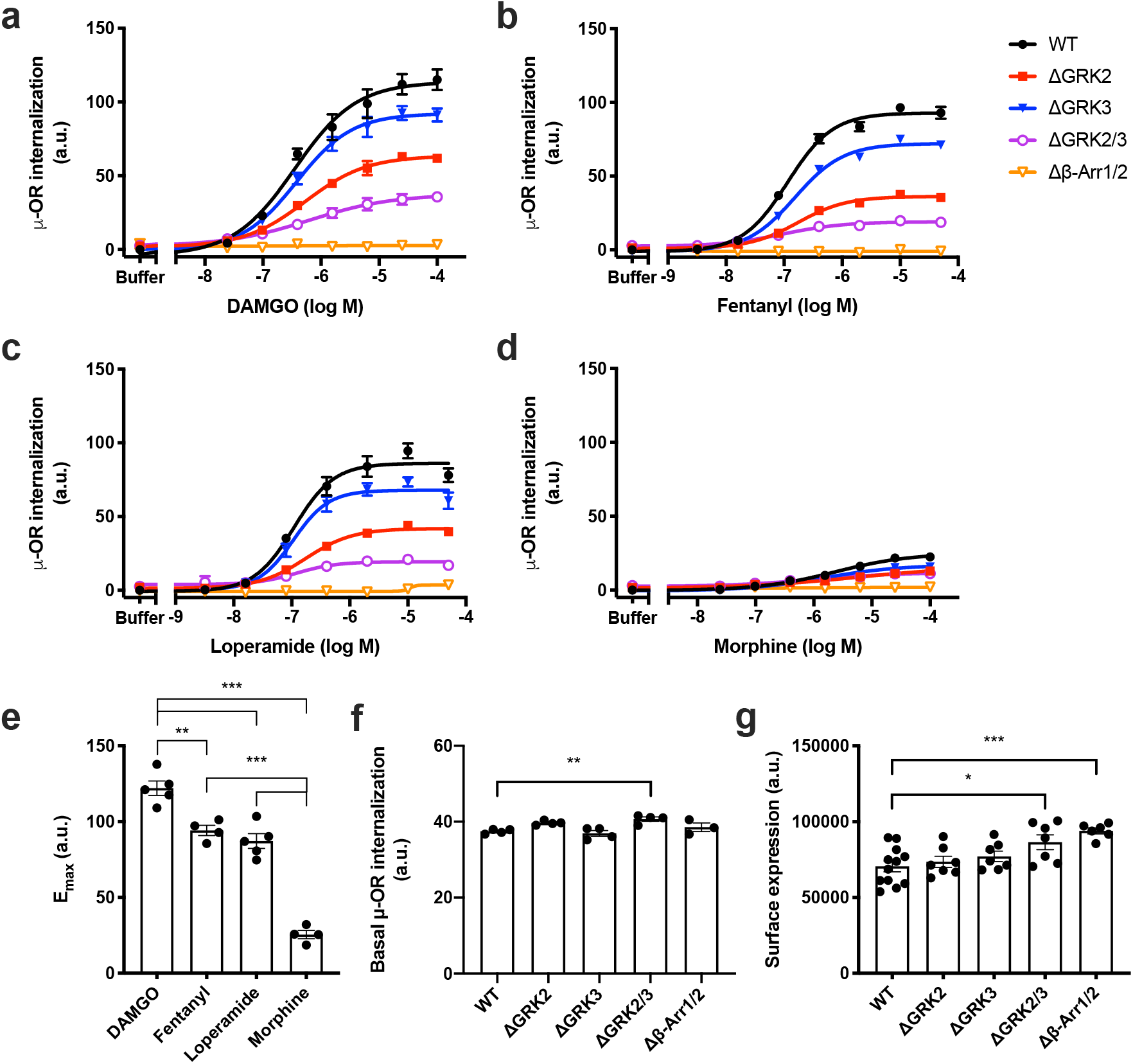
Agonist-induced internalization of μ-OR in genetically modified cell lines. Internalization of μ-OR in response to a range of concentrations of (**a**) DAMGO, (**b**), fentanyl (**c**) loperamide, or (**d**) morphine in parental HEK293A (WT) cells or HEK293A cell lines with deletion of GRK2 (ΔGRK2), GRK3 (ΔGRK3), GRK2 and −3 (ΔGRK2/3) or β-arrestin1 and −2 (Δβ-Arr1/2). Data represent the mean ± SEM of the area under the curve of 90 min real-time internalization experiments after subtraction of the buffer response in parental cells from 3-5 independent experiments carried out in duplicate. Error bars not shown lie within the dimension of the symbol. (**e**) E_max_ of agonist responses in parental cells. Statistical comparison by one-way ANOVA with Dunnett’s multiple comparisons test. (**f**) Basal μ-OR internalization determined by taking the area under the curve of the buffer responses. (**g**) μ-OR cell surface expression determined by measuring donor signals in absence of acceptor. (**e-g**) Data represent the mean of individual experiments (circles) as well as the mean ± SEM (columns) from 4-5 (**e**), 3-4 (**f**) or 6-9 (**g**) independent experiments with 2 (**e**), 4-8 (**f**) or 32 (**g**) replicates per experiment. (**f-g**) Values in knockout cells were compared to the parental cells by one-way ANOVA with Dunnett’s multiple comparisons test. \**P* = 0.01-0.05, \*\**P* = 0.001-0.01, \*\*\**P* < 0.001. a.u., arbitrary units.

Internalization was clearly reduced in the ΔGRK cells stimulated with DAMGO, loperamide and fentanyl. A similar tendency was observed for morphine, but the effect was less clear compared to the three other agonists due to lower overall internalization levels upon morphine stimulation. A cell line where β-arrestin1 and −2 have been deleted in the same cellular background as the ΔGRK cell lines has previously been described^35^. None of the agonists were able to induce internalization in the Δβ-arrestin1/2 cell line (**Fig. 2, Supplementary Fig. S3**). We used the area under the curve from the 90 min real-time internalization ratio traces to plot concentration-response curves and determine the maximum response (E_max_) and potency (EC_50_) for the four agonists in each of the cell lines where internalization could be detected (**Fig. 2, Table 1**). DAMGO stimulation led to a higher E_max_ in the parental cells than stimulation with fentanyl and loperamide, which was larger than the E_max_ upon morphine stimulation (**Fig. 2e**). For DAMGO, fentanyl and loperamide, we found a decrease in E_max_ compared to the parental cells of 48-62% and 22-23% in the ΔGRK2 and the ΔGRK3 cell lines, respectively. The E_max_ was further reduced, but not completely abolished in the ΔGRK2/3 cell line and corresponded approximately to the sum of the reductions in the individual ΔGRK2 and ΔGRK3 cell lines. The responses to morphine stimulation were too small for in-depth quantitative analysis. Importantly, the μ-OR surface expression in the ΔGRK2 and/or −3 cell lines was similar or slightly higher than in the parental cells (**Fig. 2g**). More DNA was required for transfection of the ΔGRK3 (30 ng/ml) and the ΔGRK2/3 (20 ng/ml) cell lines compared to the parental and the ΔGRK2 (15 ng/ml) cell lines to achieve this. To determine whether the reduced internalization was due to off-target effects of the single guide RNAs (sgRNAs) we overexpressed the deleted GRKs in the corresponding cell lines. GRK2 or −3 overexpression restored the internalization ratio to the levels of the parental cells, thus showing that the internalization machinery is fully functional in the ΔGRK cell lines (**Supplementary Fig. S4**).

**Table 1.**
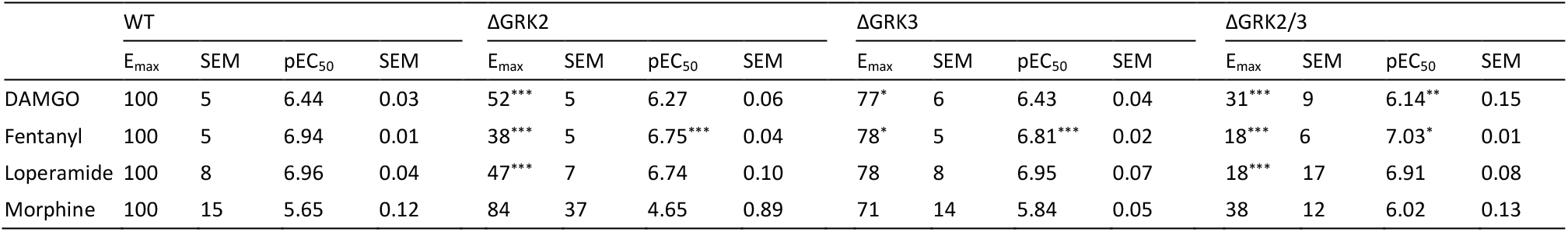
E_max_ (% of parental HEK293A (WT) cells) and pEC_50_ values for μ-OR internalization obtained from fitting concentration-response curves calculated by taking the area under the curve of 90 min internalization time curves to a four-parameter model of agonism. No fit could be obtained for experiments in cells where β-arrestin1 and −2 had been deleted. E_max_ and pEC_50_ values are the mean of 3-5 independent experiments. E_max_ and pEC_50_ in knockout cells are compared to the parental cells by one-way ANOVA with Dunnett’s multiple comparisons test. \**P* = 0.01-0.05, \*\**P* = 0.001-0.01, \*\*\**P* < 0.001.

| | WT | | | | ΔGRK2 | | | | ΔGRK3 | | | | ΔGRK2/3 | | | |
| --- | --- | --- | --- | --- | --- | --- | --- | --- | --- | --- | --- | --- | --- | --- | --- | --- |
| | $E_{\max}$ | SEM | $pEC_{50}$ | SEM | $E_{\max}$ | SEM | $pEC_{50}$ | SEM | $E_{\max}$ | SEM | $pEC_{50}$ | SEM | $E_{\max}$ | SEM | $pEC_{50}$ | SEM |
| DAMGO | 100 | 5 | 6.44 | 0.03 | 52*** | 5 | 6.27 | 0.06 | 77* | 6 | 6.43 | 0.04 | 31*** | 9 | 6.14** | 0.15 |
| Fentanyl | 100 | 5 | 6.94 | 0.01 | 38*** | 5 | 6.75*** | 0.04 | 78* | 5 | 6.81*** | 0.02 | 18*** | 6 | 7.03* | 0.01 |
| Loperamide | 100 | 8 | 6.96 | 0.04 | 47*** | 7 | 6.74 | 0.10 | 78 | 8 | 6.95 | 0.07 | 18*** | 17 | 6.91 | 0.08 |
| Morphine | 100 | 15 | 5.65 | 0.12 | 84 | 37 | 4.65 | 0.89 | 71 | 14 | 5.84 | 0.05 | 38 | 12 | 6.02 | 0.13 |

### GRK2 and GRK3 deletion reduces β-arrestin2 recruitment

To further investigate the mechanism linking GRK2/3 and μ-OR internalization we measured the recruitment of β-arrestin2 to μ-OR in the ΔGRK2 and/or −3 cells using a bioluminescence resonance energy transfer 1 (BRET^1^) assay. In this assay, BRET between *Renilla reniformis* luciferase II (RlucII)-tagged β-arrestin2 and μ-OR tagged with EYFP at the C-terminus is measured in real-time. DAMGO, fentanyl and loperamide stimulation induced a peak in β-arrestin2 recruitment at 6 min for DAMGO and at 6-12 min for fentanyl and loperamide followed by a gradual decrease in recruitment in the parental and ΔGRK3 cells (**Fig. 3**). The responses were strongly decreased in the ΔGRK2 and ΔGRK2/3 cell lines and in most cases too small to discern the kinetic profiles, except in the ΔGRK2 cells with DAMGO stimulation where a similar time-course was observed. The response for morphine was too low to detect a peak in the β-arrestin2 recruitment. Since the kinetic profile was similar for all conditions where the response was sufficiently large, we used the 6 min peak response to generate concentration-response curves for determination of the maximum responses (E_max_) and EC_50_ values for the β-arrestin2 recruitment (**Supplementary Fig. S5, Supplementary Table S1**). In the parental HEK293A cells, DAMGO stimulation led to the highest maximum response (E_max_), followed by loperamide, fentanyl and morphine stimulation. The E_max_ in the ΔGRK3 cells was similar to the parental cells, but reduced to 19-25% of the E_max_ in the parental cells in the ΔGRK2 cells stimulated with DAMGO and loperamide. The β-arrestin2 recruitment in the ΔGRK2 cells stimulated with fentanyl and morphine and in the ΔGRK2/3 cells stimulated with all ligands was too low for robust curve fitting and in most cases no increase was seen even at the highest ligand concentrations. In the absence of agonist stimulation, the β-arrestin2 recruitment was similar in all four cell lines (**Supplementary Fig. S5**).

**Figure 3.**
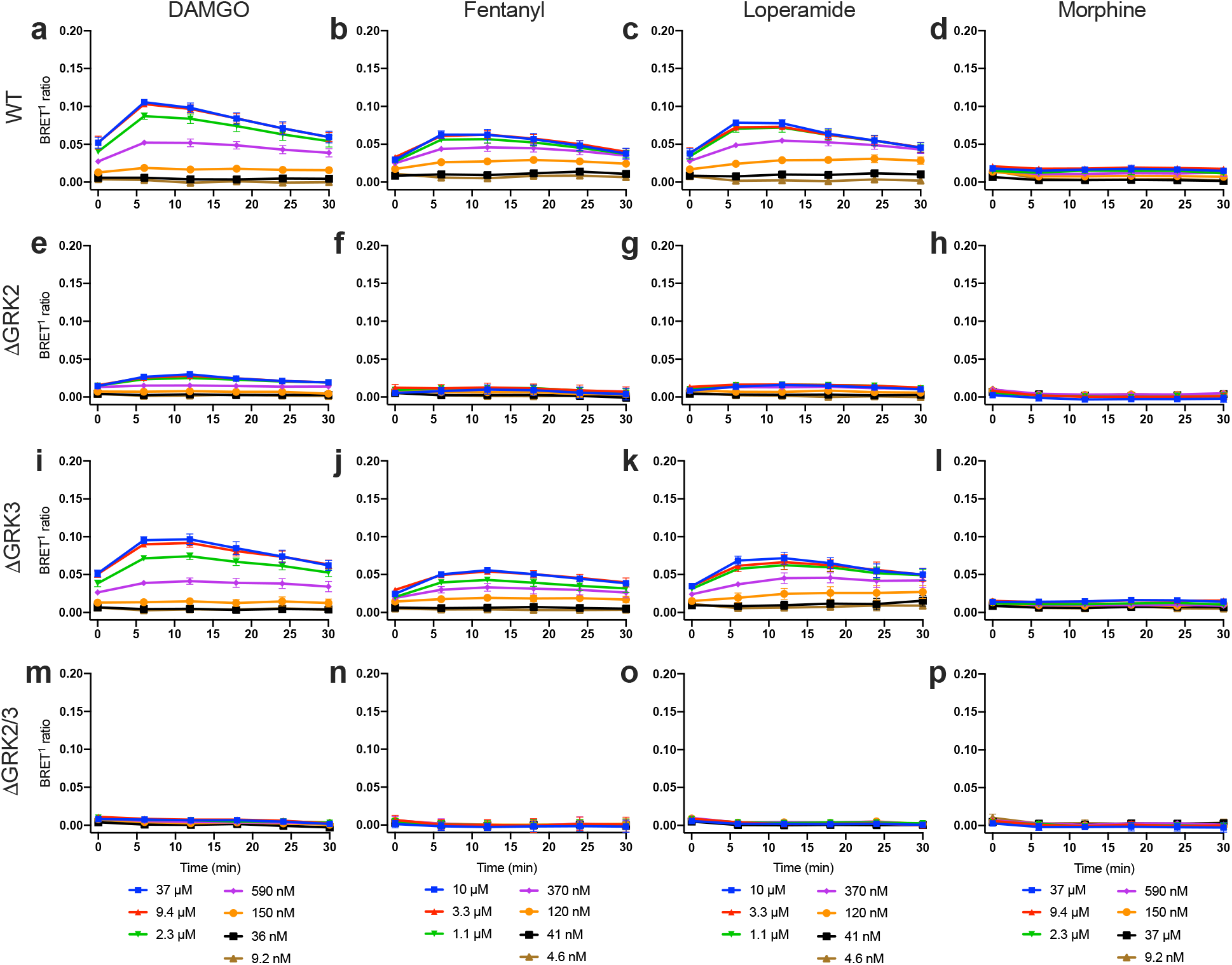
Time course of β-arrestin2 recruitment to μ-OR in GRK knockout cell lines. Recruitment of β-arrestin2-RlucII to μ-OR-EYFP in response to stimulation of μ-OR by a range of concentrations of DAMGO, fentanyl, loperamide or morphine in parental (WT) cells (**a-d**), ΔGRK2 cells (**e-h**), ΔGRK3 cells (**i-l**) or ΔGRK2/3 cells (**m-p**). Data represent mean ± SEM of the BRET^1^ ratio after buffer subtraction from 3 independent experiments carried out in duplicate. Error bars not shown lie within the dimension of the symbol.

We further investigated the β-arrestin2 recruitment to μ-OR using an enhanced bystander BRET (ebBRET) assay^36^. Here, BRET between β-arrestin2-RlucII and a membrane anchored *R. reniformis* GFP (rGFP) increases as β-arrestin2 is recruited to the membrane. Importantly, the C-terminus of the receptor is not modified for this assay and the amount of μ-OR DNA used for transfection could be reduced by around 20-fold compared to the μ-OR-EYFP assay described above while maintaining a good assay window. In agreement with the μ-OR-EYFP assay, β-arrestin2 recruitment was similar in the parental and the ΔGRK3 cells and clearly decreased in the ΔGRK2 and ΔGRK2/3 cell lines after 6 min stimulation with DAMGO, fentanyl and loperamide (**Fig. 4**). The maximum responses (E_max_) in the ΔGRK2 and ΔGRK2/3 cells were 51-59% and 38-42%, respectively, of the E_max_ in the parental cells (**Table 2**). These values are comparable to the corresponding E_max_ values in the internalization assay, but higher than the μ-OR-EYFP β-arrestin2 recruitment assay. Increasing the amount of μ-OR DNA used for transfection in the ebBRET assay to an amount comparable to the μ-OR-EYFP assay did not reduce the responses in the ΔGRK2 and ΔGRK2/3 cell lines (**Supplementary Fig. S6**). Rescue expression of either GRK2 or GRK3 in the three ΔGRK cell lines increased β-arrestin2 recruitment to the level of the parental cells with overexpression of the same GRK for all four ligands (**Supplementary Fig. S7**).

**Table 2.** E_max_ (% of parental HEK293A (WT) cells) and pEC_50_ values for β-arrestin2-RlucII recruitment to the plasma membrane measured by ebBRET after 6 min of μ-OR stimulation. The values are obtained from fitting concentration-response curves to a four-parameter model of agonism. E_max_ and pEC_50_ values are the mean of 3-4 independent experiments. E_max_ and pEC_50_ in knockout cells are compared to the parental cells by one-way ANOVA with Dunnett’s multiple comparisons test. \**P* = 0.01-0.05, \*\**P* = 0.001-0.01.

| | WT | | | | ΔGRK2 | | | | ΔGRK3 | | | | ΔGRK2/3 | | | |
| --- | --- | --- | --- | --- | --- | --- | --- | --- | --- | --- | --- | --- | --- | --- | --- | --- |
| | $E_{\max}$ | SEM | $pEC_{50}$ | SEM | $E_{\max}$ | SEM | $pEC_{50}$ | SEM | $E_{\max}$ | SEM | $pEC_{50}$ | SEM | $E_{\max}$ | SEM | $pEC_{50}$ | SEM |
| DAMGO | 100 | 8 | 6.56 | 0.05 | 59* | 9 | 6.41 | 0.15 | 107 | 10 | 6.4 | 0.05 | 38** | 8 | 6.56 | 0.09 |
| Fentanyl | 100 | 9 | 6.99 | 0.1 | 57* | 7 | 6.78 | 0.16 | 107 | 10 | 6.93 | 0.06 | 38** | 13 | 7.32 | 0.12 |
| Loperamide | 100 | 14 | 6.95 | 0.11 | 51 | 16 | 6.76 | 0.28 | 102 | 12 | 6.77 | 0.02 | 42* | 13 | 6.65 | 0.36 |
| Morphine | 100 | 13 | 5.78 | 0.35 | 69 | 46 | n.d. |  | 97 | 23 | n.d. |  | 74 | 27 | n.d. |  |

**Figure 4.**
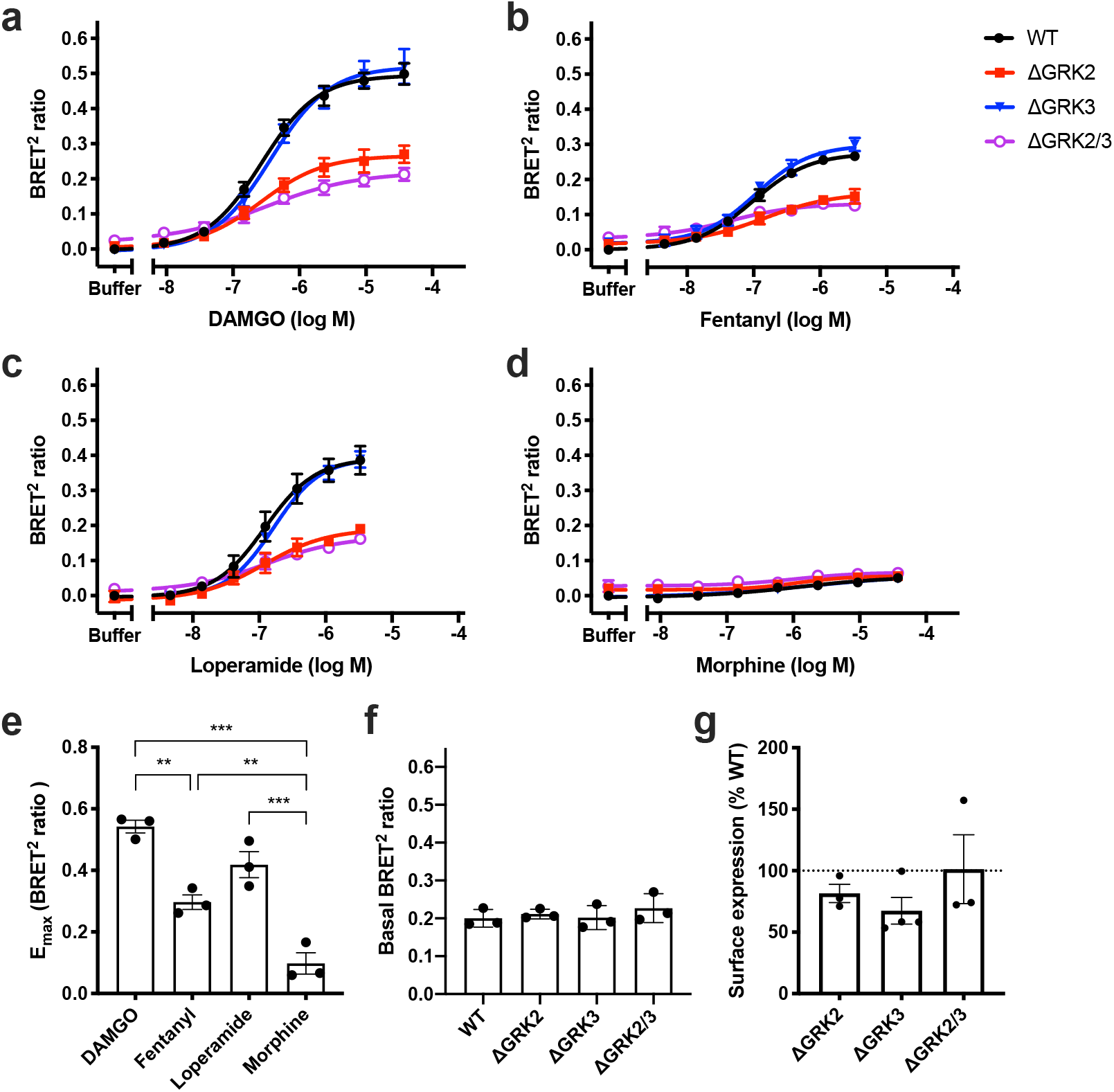
β-arrestin2 recruitment of μ-OR in genetically modified cell lines. Recruitment of β-arrestin2-RlucII to the cell membrane monitored by ebBRET in response to 6 min stimulation of μ- OR by a range of concentrations of (**a**) DAMGO, (**b**) fentanyl, (**c**) loperamide, or (**d**) morphine in parental HEK293A (WT) cells or HEK293A cell lines with deletion of GRK2 (ΔGRK2), GRK3 (ΔGRK3), or GRK2 and −3 (ΔGRK2/3). Data represent the mean ± SEM of the BRET^2^ ratio after subtraction of the buffer response in parental cells from 3-4 independent experiments carried out in duplicate. Error bars not shown lie within the dimension of the symbol. (**e**) E_max_ of agonist responses in parental cells. Statistical comparison by one-way ANOVA with Dunnett’s multiple comparisons test. \*\**P* = 0.001-0.01, \*\*\**P* < 0.001. (**f**) Basal β-arrestin2 recruitment determined as the buffer response. (**g**) μ-OR surface expression measured by ELISA and normalized to expression in parental cells. (**e-g**) Data represent the mean of individual experiments (circles) and the mean ± SEM (columns) from 3 (**e-f**) or 3-4 (**g**) independent experiments with 2 (**e**), 6-8 (**f**) or 3 (**g**) replicates per experiment. (**f-g**) Values in knockout cells were compared to the parental cells by one-way ANOVA with Dunnett’s multiple comparisons test and no significant differences were found (*P* > 0.05).

### Sustained and GRK2/3 independent recruitment of β-arrestin2 to the plasma membrane

We have previously observed that a considerable amount of β-arrestin2 recruitment can be detected in the ebBRET assay after 60 min of μ-OR stimulation^25^. To determine if there is a kinetic component to the effect of GRK2 and/or −3 knockout we measured the β-arrestin2 recruitment in the ebBRET assay after 60 min stimulation in the genome-edited cell lines. There was a substantial increase in β-arrestin2 recruitment at high agonist concentrations (**Supplementary Fig. S8**), although the recruitment in the parental cells was slightly lower after 60 min than after 6 min stimulation (**Fig. 5a,b**). The E_max_ values in the parental cells followed the same distribution as observed after 6 min stimulation (**Supplementary Fig. S8**) and the EC_50_ values were similar to the values obtained after 6 min stimulation (**Supplementary Table S2**). However, β-arrestin2 recruitment was unchanged in the ΔGRK2 and/or −3 cell lines relative to the parental cells, with the exception of the ΔGRK3 cell line where DAMGO stimulation led to a 40% increased recruitment. In contrast, the agonist-induced β-arrestin2 recruitment to μ-OR-EYFP was almost reduced to buffer levels after 60 min stimulation (**Fig. 5c,d**). To get further insights into the mechanism behind this difference we used a third β-arrestin2 recruitment assay where recruitment of GFP^2^-β-arrestin2 (BRET acceptor) to μ-OR-RlucII (BRET donor) is measured by BRET^2^. The results of this assay were overall in good agreement with the μ-OR-EYFP assay (**Fig. 5e,f**). Recruitment of GFP^2^-β-arrestin1 to μ-OR-RlucII yielded results similar to recruitment of GFP^2^-β-arrestin2 (**Supplementary Fig. S9**).

**Figure 5.**
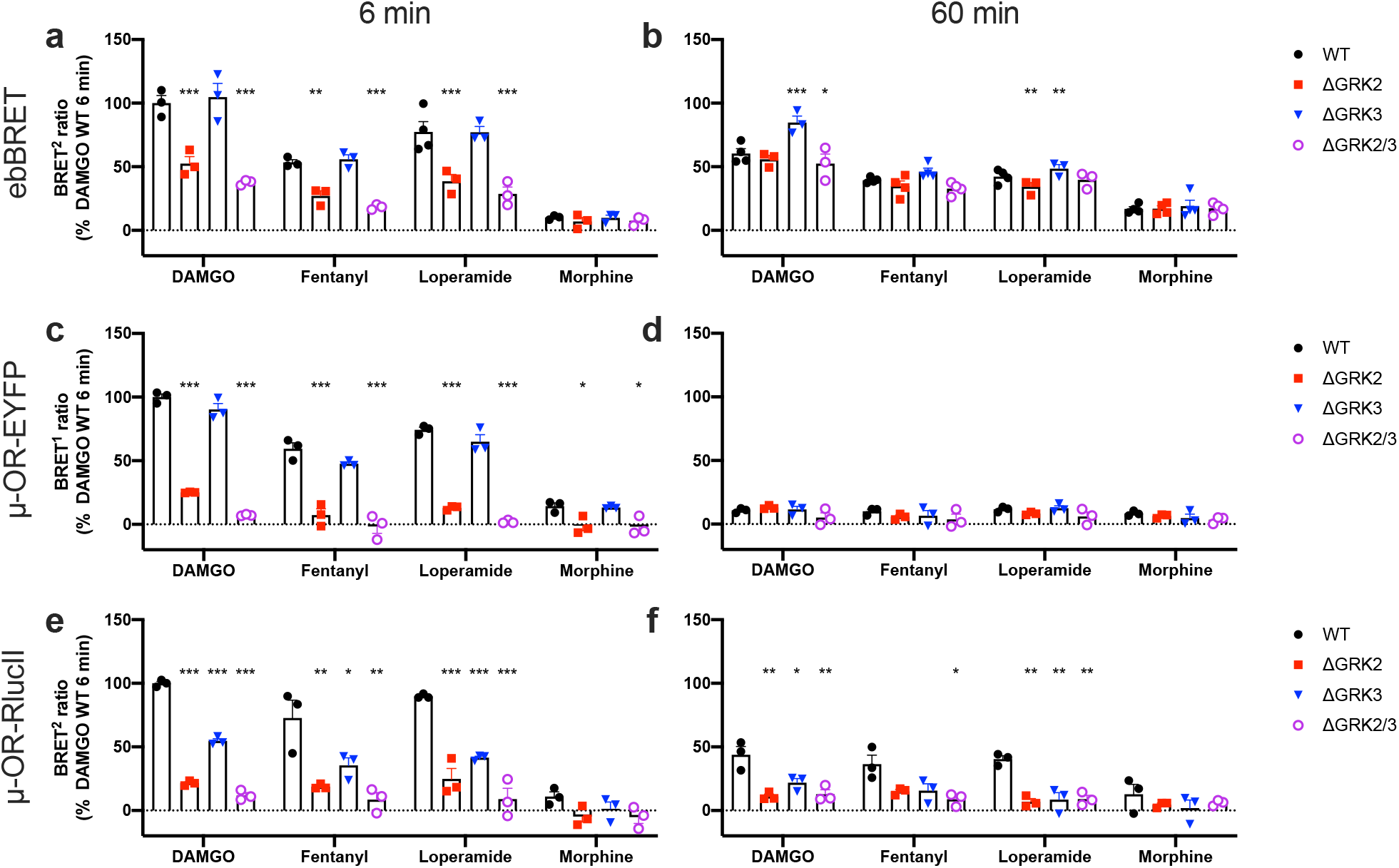
Time dependence of β-arrestin2 recruitment of μ-OR in ΔGRK cell lines. Recruitment of β-arrestin2 after 6 min (**a**, **c**, **e**) or 60 min (**b**, **d**, **f**) stimulation by saturating concentrations of DAMGO (37.5 μM), fentanyl [3.3 μM (**a-b**) or 10 μM (**c-f**)], loperamide [3.3 μM (**a-b**) or 10 μM (**c-f**)] or morphine (37.5 μM) measured by three different assays: ebBRET (**a-b**), recruitment of β-arrestin2-RlucII to μ-OR-EYFP (**c-d**), or recruitment of GFP^2^-β-arrestin2 to μ-OR-RlucII. Data represent the mean of individual experiments (circles) as well as the mean ± SEM (columns) after buffer subtraction and normalization to the maximum response in parental (WT) cells stimulated with DAMGO for 6 min from 3-4 independent experiments performed in duplicate. BRET ratios in knockout cells were compared with parental cells by repeated measures one-way ANOVA with Dunnett’s multiple comparisons test, except for ebBRET assay with 60 min loperamide stimulation, which was compared by mixed-effects analysis with Dunnett’s multiple comparisons test. \**P* = 0.01-0.05, \*\**P* = 0.001-0.01, \*\*\**P* < 0.001.

Finally, we looked at the kinetic profile of the effect of GRK deletion on μ-OR internalization. We initially analyzed the μ-OR internalization by taking the area under the curve of the 90 min time curves, which could mask kinetic effects. However, concentration-response curves composed from the μ-OR internalization responses after 6, 12, 30 and 60 min stimulation were highly similar for all curves displaying sufficient signal-to-noise (**Supplementary Fig. S10**), thus indicating that the effect of GRK2/3 knockout on μ-OR internalization is constant over time.

### Similar effects of genetic deletion and pharmacological inhibition of GRK2/3

The small molecule inhibitor CMPD101 has been shown to selectively inhibit GRK2/3 over GRK1 and GRK5 *in vitro*^30^. We compared the internalization and β-arrestin2 recruitment results obtained in the ΔGRK2/3 cells with that obtained using the kinase inhibitor. We used the ebBRET assay with 6 min stimulation to measure the β-arrestin2 recruitment. At CMPD101 concentrations ≤ 10 μM, we observed no effect on receptor internalization in absence of agonist (basal internalization) (**Fig. 6a**) or basal β-arrestin2 recruitment (**Fig. 6e**) in the parental HEK293A and ΔGRK2/3 cells. With DAMGO stimulation in the parental cells, we found a concentration dependent decrease in μ-OR internalization (**Fig. 6b**) and β-arrestin2 recruitment (**Fig. 6f**) that plateaued at levels similar to the ΔGRK2/3 cells with IC_50_ values of 1.8 μM (pIC_50_ = 5.79 ± 0.12) for internalization and 0.95 μM (pIC_50_ = 6.03 ± 0.02) for β-arrestin2 recruitment at CMPD101 concentrations ɤ 10 μM. Thus, in this concentration range CMPD101 specifically inhibits GRK2/3 and reaches inhibition comparable to the ΔGRK2/3 cell line at 10 μM.

**Figure 6.**
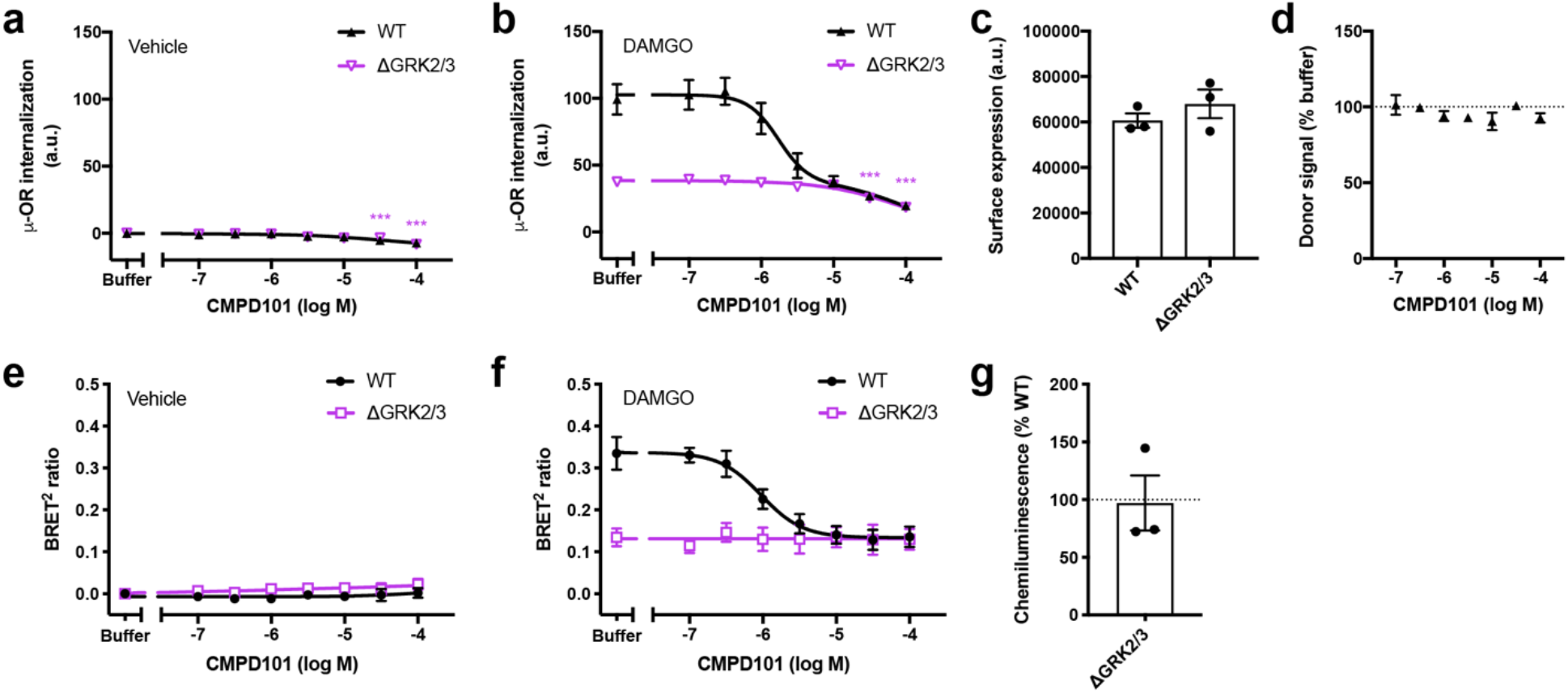
GRK2/3 inhibition and off-target effects by CMPD101. Effect of a range of CMPD101 concentrations on (**a**) basal (vehicle treated) or (**b**) 100 μM DAMGO induced μ-OR internalization and (**e**) basal (vehicle treated) or (**f**) 37.5 μM DAMGO stimulated β-arrestin2 recruitment in parental HEK293A (WT) cells or HEK293A cell lines with deletion of GRK2 and −3 (ΔGRK2/3). β-arrestin2 recruitment was measured by ebBRET after 6 min stimulation of μ-OR. Data represent the mean ± SEM after buffer (vehicle treated) or vehicle (DAMGO treated) subtraction from 3 independent experiments carried out in duplicate. μ-OR surface expression was determined from donor signals in absence of acceptor for internalization experiments (**c**) and by ELISA for β-arrestin2 recruitment experiments (**g**). Data points represent the mean of individual experiments and bars represent the mean ± SEM from three independent experiments with 32 (internalization) or 3 (ELISA) replicates per experiment. In (**g**) data were normalized to expression in parental cells. (**d**) Effect of a range of CMPD101 concentrations on donor signals in absence of acceptor. An unaffected donor signal indicates that the compound does not interfere with the assay. Data represent the mean ± SEM from 3 independent experiments carried out in duplicate. Effect of different CMPD101 concentrations on ΔGRK2/3 cells in (**a**), (**b**), (**d**), (**e**) and (**f**) was compared to buffer by one-way repeated measures ANOVA with Dunnett’s multiple comparisons test before normalization. Expression in parental and ΔGRK2/3 cells in (**c**) and (**g**) was compared by ratio paired t-test before normalization. \*\*\**P* < 0.001. a.u., arbitrary units.

CMPD101 concentrations ≥ 30 μM decreased basal and DAMGO-stimulated μ-OR internalization in both parental and ΔGRK2/3 cells (**Fig. 6a,b**). In contrast, basal and DAMGO induced β-arrestin2 recruitment were constant over the range of CMPD101 concentrations tested (up to 100 μM) in parental and ΔGRK2/3 cells (**Fig. 6e,f**). μ-OR surface expression was similar in the parental and ΔGRK2/3 cells in the internalization and β-arrestin2 recruitment experiments (**Fig. 6c,g**). Due to the long-lifetime donor fluorophore with several emission peaks and the homogenous format, the internalization assay is sensitive to compounds that absorb light in most of the visual spectrum. However, the donor signal in absence of acceptor was unaffected by CMPD101 up to concentrations of 100 μM (**Fig. 6d**), demonstrating that the observed effects at high concentrations were not due to optical interference with the internalization assay. We conclude that CMPD101 at concentrations ≥ 30 μM has a non-GRK2/3 regulatory effect on basal and agonist-stimulated μ-OR internalization.

## Discussion

Compelling evidence on the importance of GRKs in μ-OR pharmacology has increased over the years. Yet, most of these studies has relied on methods either overexpressing dominant negative forms of the protein of interest or silencing with siRNAs, possibly leading to undesired effects of overexpression or incomplete removal of the target protein^37^. Genome editing is a promising alternative to these approaches to investigate the role of GRKs as potential modulators of GPCR function. In this study we have generated genome-edited HEK293 cells and used them to dissect the role of GRK2 and GRK3 in μ-OR internalization and β-arrestin2 recruitment.

Previous studies have demonstrated that GRK2 and GRK3 affect the phosphorylation state of the ^375^STANT^379^ motif in mouse μ-OR^22,23,26^, and this region has been directly linked to the regulation of μ-OR internalization by mutational studies^23,24^. Here, we demonstrate that our ΔGRK2, ΔGRK3 and ΔGRK2/3 cell lines are excellent novel tools to dissect the role of these GRK subtypes in DAMGO, fentanyl and loperamide-induced μ-OR internalization. Double KO of GRK2/3 had an additive effect on μ-OR internalization with a significant reduction for DAMGO-, fentanyl-and loperamide-induced μ-OR internalization (**Fig. 2**, **Table 1**). No statistically significant difference could be reached for morphine-induced μ-OR internalization, however this agonist induced low μ- OR internalization and thereby the effects could be masked by the small assay window. Our study also shows that there are additional factors controlling μ-OR internalization, since deletion of GRK2/3 did not fully inhibit μ-OR internalization. There is limited evidence suggesting that GRK5 or GRK6 could be responsible for this: μ-OR internalization is sensitive to GRK5 and GRK6 overexpression^16,25^ and GRK5 has been shown to be important in HEK293 cells and mouse brain for morphine-induced phosphorylation of Ser375^26^, which is a critical residue in initiating further C-terminal μ-OR phosphorylation^23^. In contrast, DAMGO-induced phosphorylation of the C-terminal region of μ-OR was shown to be unaffected by knockdown of GRK5 and GRK6^23,26^ and fentanyl-induced Ser375 phosphorylation was unchanged in mice lacking GRK5^15^, so the roles of GRK5 and GRK6 might depend on the GRK expression levels in an individual cell. Several other kinases and enzymes have been proposed to regulate μ-OR internalization, including phospholipase D2 (PLD2) and protein kinase C (PKC)^5^, but more studies are required to determine their role relative to the GRKs. Our successful generation of the ΔGRK2, ΔGRK3 and ΔGRK2/3 cell lines demonstrate the utility in this approach and calls for generation of additional cell lines deficient of other kinases to dissect other contributors to μ-OR internalization in the future.

We observed a discrepancy between the two assays where β-arrestin2 recruitment is measured using an EYFP or RlucII tag on the C-terminal end of the receptor and the ebBRET assay where β-arrestin2 recruitment is measured using a BRET acceptor tethered to the membrane (**Fig. 5**). The two main differences between the assays is the presence or absence of a C-terminal tag on μ-OR and the measurement of β-arrestin2 proximity to μ-OR or the cell membrane. Although C-terminal tagging of μ-OR has previously been shown not to affect the μ-OR signaling in HEK293 cells or the behavior of knock-in mice^27^ and we obtained similar results with two different tags, it cannot be ruled out that the tags could interfere with the GRK and/or β-arrestin interaction with μ-OR. Another potential explanation for the observed sustained β-arrestin recruitment to the cell membrane is that there is a GRK2/3 independent component in addition to the GRK2/3 dependent β-arrestin recruitment. This GRK2/3 independent component is only transiently associated with μ- OR and remains associated with the cell membrane after dissociation from μ-OR. Such a mechanism has previously been shown for several receptors, including μ-OR^38–40^, and has been suggested to result from an interaction where arrestin only engages the core of the receptor 7TM bundle^38,40–42^. This interaction model would be consistent with our observation that the β-arrestin population that remains associated with the cell membrane interacts with μ-OR independently of GRK2/3. Our results would suggest that the combination of direct and bystander β-arrestin recruitment assays that measure receptor and cell membrane proximity at different time points could be used to discriminate transient and stable β-arrestin interactions.

Although using CRISPR/Cas9 genome editing to generate cell lines lacking specific proteins has several advantages over conventional methods for interrogating protein function, it has also been criticized for potentially selecting clones that have compensated for the lack of the deleted protein^43^. We addressed this concern by reintroduction of the deleted proteins (**Supplementary Figs. S4**, **S7**) and by pharmacological inhibition with CMPD101 (**Fig. 6**). Transient expression of GRK2 or GRK3 showed that μ-OR in the genome-edited cells was still sensitive to regulation by GRK2 and GRK3. Comparison with pharmacological inhibition is an alternative way to characterize genome-edited cells if a specific and potent inhibitor exists. We compared the μ-OR internalization and β-arrestin2 recruitment in the ΔGRK2/3 cells with the effect of the inhibitor CMPD101 on the parental cells and found highly similar results. Collectively, these results demonstrate that there are no significant compensatory mechanisms in the ΔGRK2/3 cells of relevance to the μ-OR functions studied here and confirms the utility of CMPD101 as a GRK2/3 selective pharmacological tool in concentrations up to 10 μM. Our ΔGRK2/3 cells also proved to be a powerful tool to study specificity of GRK inhibitors as we could demonstrate off-target effects of CMPD101 at concentrations ≥ 30 μM (**Fig. 6a,b**), which is a concentration range commonly used to inhibit GRK2/3^10,31^, thus cautioning the use of such high concentrations in future studies.

In conclusion, we have generated novel ΔGRK2, ΔGRK3 and ΔGRK2/3 cell lines in the highly utilized HEK293A cell background. These cell lines complement the range of G protein and β-arrestin cell lines generated by Dr. Asuka Inoue^44^ and thus expand this highly efficient tool box to study intracellular proteins involved in GPCR function and signaling. Here, we demonstrate the utility of the cell lines to study μ-OR-mediated β-arrestin2 recruitment to the cell membrane and μ-OR internalization, but we envision that the cell lines could be used for a range of other studies and thus welcome all requests to obtain the ΔGRK2, ΔGRK3 and ΔGRK2/3 cell lines for future studies.

## Methods

### Materials

Dulbeccos’s modified Eagle medium (DMEM, Cat# 61965-026), fetal bovine serum (FBS), Opti-MEM, Penicillin-Streptomycin, Dulbecco’s phosphate buffered saline (DPBS, Cat# 14190-144), Hank’s balanced salt solution (HBSS, Cat# 14175-053), restriction endonucleases, TOPO TA cloning kit for sequencing including TOP10 *E. coli.*, FastAP alkaline phosphatase, Pierce BCA Protein Assay Kit, dithiothreitol, PageRuler Plus Prestained Protein Ladder (10 to 250 kDa), Pierce 10x Tris-Glycine SDS Buffer, NuPAGE LDS Sample buffer (4x), SuperSignal West Pico PLUS and ELISA Femto chemiluminescent substrates, and Pluronic F-68 non-ionic surfactant were purchased from Thermo Fisher Scientific (Waltham, MA, USA). QuickExtract DNA Extraction Solution was purchased from Lucigen Corporation (WI, USA) and TEMPase Hot Start DNA Polymerase from Ampliqon (Odense, Denmark). Anti-GRK2 antibody (Cat# MAB43391, RRID:AB_2818985) and anti-GRK5 antibody (Cat# AF4539, RRID:AB_2248068) were obtained from R&D Systems (Minneapolis, MN, USA). Anti-GRK3 antibody (Cat# 80362, RRID:AB_2799951), anti-GRK6 antibody (Cat# 5878, RRID:AB_11179210), and anti-rabbit IgG antibody conjugated to horseradish peroxidase (HRP) (Cat# 7074, RRID:AB_2099233) were acquired from Cell Signaling Technology (Danvers, MA, USA). Anti-GAPDH (Cat# NB600-502, RRID:AB_10077682) antibody was bought from Novus Biologicals (Centennial, CO, USA). Anti-mouse IgG antibody conjugated to HRP (Cat# P0447, RRID:AB_2617137) and anti-goat IgG antibody conjugated to HRP (Cat# P0449, RRID:AB_2617143) were obtained from Agilent Technologies (Santa Clara, CA, USA). Mini-PROTEAN TDX Precast Protein Gels and Trans-Blot Turbo RTA Mini PVDF Transfer Kit were purchased from Bio-Rad Laboratories (Hercules, CA, USA). T4 PNK and T4 ligase were purchased from New England Biolabs (Ipswich, MA, USA). Plasmid-Safe ATP-Dependent DNase was purchased from Epicentre Technologies Corp. (Madison, WI, USA). Primers and sgRNA sequences were purchased from TAG Copenhagen (Copenhagen, Denmark). FuGene6 Transfection Reagent was purchased from Promega (Madison, WI, USA). Polyethylenimine (PEI) was purchased from Polysciences Inc. (Warrington, PA, USA). Coelenterazine 400a and coelenterazine h were purchased from Cayman Chemical Company (Ann Arbor, MI, USA). Tag-lite SNAP Lumi4-Tb was purchased from Cisbio (Codolet, France). DAMGO was purchased from Abcam (Cambridge, United Kingdom). Anti-FLAG M2 antibody (Cat# F3165, RRID:AB_259529), morphine sulfate, fentanyl citrate, and loperamide hydrochloride, RIPA buffer, cOmplete protease inhibitor cocktail, Cell Dissociation Solution, protease inhibitor cocktail, Trizma base, skim milk powder, and Tween 20 were purchased from Sigma-Aldrich (St. Louis, MO, USA). CMPD101 was purchased from Tocris Bioscience (Bristol, UK).

### Plasmids

The following plasmids were described previously: pcDNA5/FRT/TO-FLAG-SNAP-μ-OR^25^, pcDNA3.1(+)-β-arrestin1 and pcDNA3.1(+)-β-arrestin2^45^, pcDNA3.1/Zeo-β-arrestin1-RlucII^46^, pcDNA3.1/Zeo-β-arrestin2-RlucII^47^, pcDNA3.1(+)-GFP^2^-β-arrestin2^48^ and pcDNA3.1(+)-rGFP-CAAX^36^. pSpCas9(BB)-2A-GFP (PX458) was a gift from Feng Zhang (Addgene plasmid # 48138; http://n2t.net/addgene:48138; RRID:Addgene_48138) and was described previously^49^. The pcDNA3.1-μ-OR-EYFP plasmid was a gift from the ARPEGE platform (Institut de Génomique Fonctionnelle, Montpellier, France). pcDNA3.1/Hygro(+)-SP-FLAG-μ-OR-RlucII was constructed by PCR amplification of human μ-OR1 tagged with a signal peptide (SP) (MKTIIALSYIFCLVFA) and a variant of the FLAG tag epitope (MDYKDDDDA) from pLVxi2P-SP-FLAG-hμ-OR^27^ and then subcloning it into the pcDNA3.1/Hygro(+)GFP10-RlucII vector^50^ using Gibson assembly (New England Biolabs) after digestion with NheI + AgeI.

The pcDNA3.1(+)-GRK2 and pcDNA3.1(+)-GRK3 constructs were kind gifts from Novo Nordisk A/S (Maaloev, Denmark). All μ-OR, GRK and β-arrestin sequences are the human sequences.

### Cell lines and culturing

The parental HEK293A and Δβ-arrestin1/2 cell lines were kind gifts from Dr. Asuka Inoue^35^. All cell lines were cultured in DMEM supplemented with 10% FBS and 100 U/ml Penicillin-Streptomycin at 37 °C and 5% CO_2_ in a humidified incubator.

### Design and cloning of sgRNAs

sgRNA sequences were identified using the WTSI Genome Editing (WGE) tool by screening exonic regions of the genes *ADRBK1* (RefSeq: NC_000011.10) or *ADRBK2* (RefSeq: NC_000022.11) encoding GRK2 and GRK3, respectively. sgRNAs with low predicted off target effects and binding sites upstream of regions encoding catalytic sites in the GRK proteins were chosen, resulting in the sgRNA sequence 5’-CTTCGACTCATACATCATGA-3’ binding in exon 4 within the *ADRBK1* gene and the sgRNA sequence 5’-ATTATTGGACGAGGAGGATT-3’ binding in exon 8 within the *ADRBK2* gene. Overhangs for cloning into a BsbI restriction site were placed in the ends of the sgRNAs. sgRNAs were prepared for cloning by incubating 100 μM of reverse complementary strands with T4 PNK in a thermocycler at 37 °C for 30 minutes followed by a 5-minute incubation at 95 °C and ramping down to 25 °C with 5 °C/min. The double stranded sgRNAs were ligated into BsbI digested pSpCas9(BB)-2A-GFP by incubating with T4 DNA ligase for 1 hour at 22 °C. To remove excess non-ligated DNA, the samples were treated with Plasmid-Safe ATP-Dependent DNase for 30 minutes at 37 °C.

### Generation and validation of CRISPR-Cas9 KO cell lines

HEK293A cells were seeded at a density of 2 × 10^5^ cells/well in a 6-well plate and incubated at 37 °C and 5% CO_2_ in a humidified incubator. Twenty-four hours later cells were transfected with 500 ng/well pSpCas9(BB)-2A-GFP encoding the sgRNAs using FuGene6 as the transfection reagent and the cells were incubated at 37 °C and 5% CO_2_ in a humidified incubator. Forty-eight hours later, the cells were harvested by trypsination and sorted based on their GFP expression with fluorescence assisted cell sorting (FACS) using a MoFlo Astrios Cell Sorter (Beckman Coulter, Brea, CA, USA). Cells were seeded in 96-well culture plates to isolate single clones and were incubated at 37 °C and 5% CO_2_ in a humidified incubator until ~70% confluent.

### IDAA

Single clones were harvested from 96-well plates by trypsination and DNA was extracted using QuickExtract followed by cell lysis (20 min at 65 °C, 10 min at 98 °C) as described previously^32^. A tri-primer PCR described previously^51^ was performed using TEMPase Hot Start DNA Polymerase. To amplify the *ADRBK1* locus, the following primers and concentrations were used: 0.05 μM forward primer (5’-AGCTGACCGGCAGCAAAATTGCCAGGCCCTTGGTGGAATTCTATG-3’), 0.5 μM reverse primer (5’-GGACATGCTCAGTGGCACTCTTC-3’) and 0.5 μM FAM forward primer (5’-6-FAM-AGCTGACCGGCAGCAAAATTG-3’). For amplification of the *ADRBK2* locus following primers and concentrations were used: 0.05 μM forward primer (5’-AGCTGACCGGCAGCAAAATTGCCTGGGGCATCTCATCCTTCAGC-3’), 0.5 μM reverse primer (5’-CGCCCGGCCTACAGCTTATTTC-3’) and 0.5 μM FAM forward primer (5’-6-FAM-AGCTGACCGGCAGCAAAATTG-3’). A touchdown thermocycling program was used with denaturation at 95 °C for 15 min followed by 15 cycles with an annealing temperature of 72 °C ramping down to 58 °C with 1 °C/cycle. Subsequent 24 cycles with 58 °C as annealing temperature was performed ending with elongation at 72 °C for 20 min. For both annealing cycles, denaturation and elongation was performed at 95 °C and 72 °C for 30 s, respectively. IDAA on the resulting PCR products was executed by COBO Technologies Aps (Copenhagen, Denmark).

### Genome sequencing

DNA from the genome-edited cell lines was extracted as described above for IDAA. The *ADRBK1* or *ADRBK2* regions targeted by the sgRNA were PCR amplified with the forward primers 5’-CCAGGCCCTTGGTGGAATTCTATG-3’ (*ADRBK1*) and 5’-CCTGGGGCATCTCATCCTTCAGC-3’ (*ADRBK2*) and the same reverse primers as used for IDAA. PCR products were cloned into the pCR4-TOPO TA vectors using the TOPO TA cloning kit and used to transform TOP10 bacteria. DNA from 5-10 single clones was sequenced for each genome-edited cell line to determine the modifications for all alleles.

### Western blot

Genome-edited HEK293A cells were incubated in 15-cm culture dishes at 37 °C and 5% CO_2_ in a humidified incubator until ~90% confluency and harvested with ice cold Cell Dissociation Solution. Cells were centrifuged in a tabletop centrifuge at 4 °C for 5 min at 500 × *g*. Whole cell lysates were prepared from pellets by resuspending in RIPA buffer containing cOmplete protease inhibitor cocktail. Cells were pulse sonicated for 30 s and incubated with end-over-end rotation at 4 °C for 60 min. Lysates were centrifuged in a tabletop centrifuge at 4 °C for 10 min at 15,000 × *g*. The supernatant was transferred to a clean microcentrifuge tube and the protein concentrations were determined using Pierce BCA Protein Assay Kit according to the manufacturer’s instructions. The absorbance of the samples was measured at 562 nm on an EnSpire Multimode Plate Reader (PerkinElmer), and the values were converted to protein concentrations by interpolation from a bovine serum albumin (BSA) standard curve. Western blot samples were prepared with 50 μg protein in NuPAGE LDS Sample Buffer supplemented with 100 μM dithiothreitol (DTT) and heated for 30 s at 50 °C. Subsequently they were incubated for 15 minutes at room temperature and electrophoresed using 4-20% polyacrylamide gels for 40 min at 200 V. Proteins were transferred onto a polyvinylidene difluoride (PVDF) membrane followed by one hour blocking with 5% skim milk in Tris buffered saline with Tween 20 (TBS-T; 10 mM Tris pH 7.4, 150 mM NaCl, 0.1% Tween 20). The PVDF membrane was incubated over night with anti-GRK2 (0.1 μg/ml in TBS-T with 5% skim milk), anti-GRK3 (1:2000 in TBS-T with 5% skim milk), anti-GRK5 (0.5 μg/ml in TBS-T with 5% skim milk), anti-GRK6 (1:1000 in TBS-T with 5% skim milk) or anti-GAPDH (1:5000 in TBS-T with 1% BSA) at 4 °C. The membranes were washed three times with TBS-T and incubated with secondary antibodies for one hour at room temperature with gentle agitation followed by three washes with TBS-T. Blots were developed with HRP substrate and imaged with a FluorChem HD2 system (ProteinSimple, San Jose, CA, USA) or iBright FL1500 (Thermo Fisher Scientific, Waltham, MA, USA).

### β-arrestin2 recruitment

Parental HEK293A cells or genome-edited cells were transfected with PEI for β-arrestin2 recruitment experiments as previously described.^36^ 500,000 cells/ml were transfected with the following combinations of plasmids: (1) ebBRET: 20 ng/ml β-arrestin2-RlucII (BRET donor), 500 ng/ml rGFP-CAAX (BRET acceptor), pcDNA5/FRT/TO-FLAG-SNAP-μ-OR and pcDNA3.1(+), (2) BRET^1^: 20 ng/ml β-arrestin2-RlucII (BRET donor), 500 ng/ml μ-OR-EYFP (BRET acceptor) and pcDNA3.1(+), and (3) BRET^2^: μ-OR-RlucII (BRET donor), 20 ng/ml GFP^2^-β-arrestin1 or −2 (BRET acceptor) and pcDNA3.1(+). The amounts of pcDNA5/FRT/TO-FLAG-SNAP-μ-OR and μ-OR-RlucII were adjusted for each cell line to equalize the expression: 15 ng/ml for parental HEK293A and ΔGRK2, 10 ng/ml (μ-OR-RlucII) or 20 ng/ml (pcDNA5/FRT/TO-FLAG-SNAP-μ-OR) for ΔGRK2/3 and 30 ng/ml for ΔGRK3. For rescue experiments, 20 ng/ml GRK2 or GRK3 was added in (1). The total DNA amount in all transfections was adjusted to 1 μg/ml with pcDNA3.1(+). 32,000 cells mixed with DNA and PEI were added to each well in poly-D-lysine-coated white, opaque CulturPlate-96 96-well plates (PerkinElmer, Waltham, MA, USA). Forty-eight hours after transfection, cells were washed and incubated in assay buffer (HBSS supplemented with 1 mM CaCl_2_, 1 mM MgCl_2_, 20 mM HEPES and 0.01% Pluronic F-68, pH 7.4) for 30 min at 37 °C before addition of agonists. For experiments with CMPD101, cells were incubated for an additional 30 min in presence of CMPD101 before agonist addition. Cells were incubated with agonists at 37 °C. For ebBRET/BRET^2^ experiments, coelenterazine 400a was added to a final concentration of 2.5 μM and 2 min later BRET was measured on an EnVision 2104 Multilabel Reader (PerkinElmer) using 410/80 nm (donor) and 515/30 nm (acceptor) emission filters. BRET^2^ ratios were calculated as the ratio of acceptor and donor emission (515 nm/410 nm). For concentration-response curves the buffer response of the parental cells was subtracted. For BRET^1^ experiments, coelenterazine h was added to a final concentration of 5 μM for 3 min before measuring BRET on an EnVision 2104 Multilabel Reader using 470/24 nm (donor) and 535/30 nm (acceptor) emission filters. BRET^1^ ratios were calculated as the ratio of acceptor and donor emission (535 nm/470 nm). For concentration-response curves the buffer response of the same cell line was subtracted.

### ELISA

Receptor surface expression in β-arrestin2 recruitment experiments was determined using enzyme-linked immunosorbent assay (ELISA) as previously described^52^ with anti-FLAG as primary antibody in a 1:1000 dilution.

### TR-FRET real-time internalization

Parental HEK293A cells or genome-edited cells were transfected with PEI for real-time internalization experiments with different amounts of pcDNA5/FRT/TO-FLAG-SNAP-μ-OR depending on the cell line (same amounts for each cell line as in β-arrestin2 recruitment experiments) to equalize expression and the total DNA amount was adjusted to 1 μg/ml with pcDNA3.1(+). For rescue experiments, cells were co-transfected with 30 ng/ml GRK2, GRK3, −β-arrestin1, or −β-arrestin2. 14,000 cells mixed with DNA and PEI were added to each well in a poly-D-lysine-coated white, opaque 384-well plates (Greiner Bio-One, Kremsmünster, Austria). Forty-eight hours after transfection, cells were labeled with 10 μl Tag-lite Lumi4-Tb for 60 min at 37 °C in Opti-MEM. After labeling, cells were washed twice with assay buffer (HBSS supplemented with 1 mM CaCl_2_, 1 mM MgCl_2_, 20 mM HEPES and 0.01% Pluronic F-68, pH 7.4) and incubated for 5 min with 10 μl of 100 μM fluorescein-O’-acetic acid. The donor signal was measured before removing the second wash from the plate and used as a measure of the μ-OR surface expression. For experiments with CMPD101, the compound was added together with fluorescein-O’-acetic acid and the incubation was extended to 30 min. 10 μl agonist was then added and internalization was read immediately after for 90 min in 6 min intervals on an EnVision 2104 Multilabel Reader using a 340/60 nm excitation filter and emission was recorded through 520/8 nm (acceptor) and 615/8.5 nm (donor) emission filters. Internalization ratios were calculated as the ratio of donor over acceptor emission (615 nm/520 nm) for real-time internalization curves. For concentration-response curves the area under the real-time internalization curves was calculated and the buffer response subtracted. The IC_50_ of CMPD101 inhibition of internalization was determined by fitting the concentration-response curve obtained by subtracting the DAMGO − buffer response in ΔGRK2/3 cells from the DAMGO − buffer response in parental cells.

### Data analysis and statistics

Data are presented as mean ± SEM of n ≥ 3 independent experiments. Results were analyzed using Prism 7 and 8 (GraphPad Software, San Diego, CA, USA). Differences were determined with regular or repeated measures one-way ANOVA with Dunnett’s multiple comparisons test or paired t-test on non-normalized data; *P* < 0.05 was considered significant. In Fig. 1b where data was compared to a reference value without variance (100 ± 0), observations were compared to the reference value using a one sample t-test and the cutoff for what was considered a significant difference was adjusted to *P* < 0.05/n (n represents the number of compared observations) to account for multiple comparisons.

## Data Availability

The datasets generated during the current study are available from the corresponding authors on reasonable request.

## Author contributions

T.C.M., M.F.P., J.M.M., M.B. and H.B.-O. participated in research design. T.C.M., M.F.P., J.R,v.S. and S.D.S. conducted experiments. T.C.M., M.F.P., J.R.v.S. and S.D.S. performed data analysis. T.C.M., M.F.P., S.D.S. and H.B.-O. wrote or contributed to drafting the manuscript. All authors edited and approved the final manuscript.

## Acknowledgements

We thank Dr. Asuka Inoue for the HEK293 parental and Δβ-arrestin1/2 cell lines and for fruitful discussions and Jens Peter Stenvang for technical assistance with cell sorting. We also thank Eric Paul Bennett for input related to indel detection by amplicon analysis. H.B.-O. acknowledges financial support from the Independent Research Fund Denmark | Medical Sciences (4183-00131A), the Lundbeck Foundation, the Novo Nordisk Foundation (NNF17OC0027004), the Carlsberg Foundation (CF17-0132) and the Toyota Foundation (9310-F). M.F.P. thank the Oticon Foundation (16-0668) and the Knud Højgaards Foundation (16-02-0142) for financial support. T.C.M. acknowledges funding from the European Union’s Horizon2020 research and innovation programme under the Marie Sklodowska-Curie grant agreement No 797497. M.B. acknowledge the financial support from a Canadian Institute for Health Research (CIHR) Foundation grant (FDN-148431). M.B. also holds a Canada Research Chair in Signal Transduction and Molecular Pharmacology.

## Additional Information

**Supplementary information** accompanies this paper at …

## Competing Interests

The authors declare no competing interests.

## Notes

### Competing Interest Statement

The authors have declared no competing interest.

### Summary of Updates

This version of the manuscript has been revised to elucidate the cause of the differences between the GRK dependence of the internalization of the mu opioid receptor and the independence of the arrestin recruitment that we reported in the original version of the manuscript. We now show that with short stimulation time in the arrestin recruitment assay there is full agreement between the GRK dependence of the internalization and the arrestin recruitment assays. In addition, with longer stimulation time we discovered by using a combination of arrestin recruitment assays that measure either receptor or plasma membrane proximity that there is a sustained, GRK independent arrestin recruitment to the plasma membrane. Furthermore, we have added western blots of the GRK5 and GRK6 expression in the knockout cells and additional controls to the rescue experiments.

