## Supplementary information for "Dissecting the roles of GRK2 and GRK3 in μ-opioid receptor internalization and β-arrestin2 recruitment using CRISPR/Cas9-edited HEK293 cells"

Supplementary Figure S1

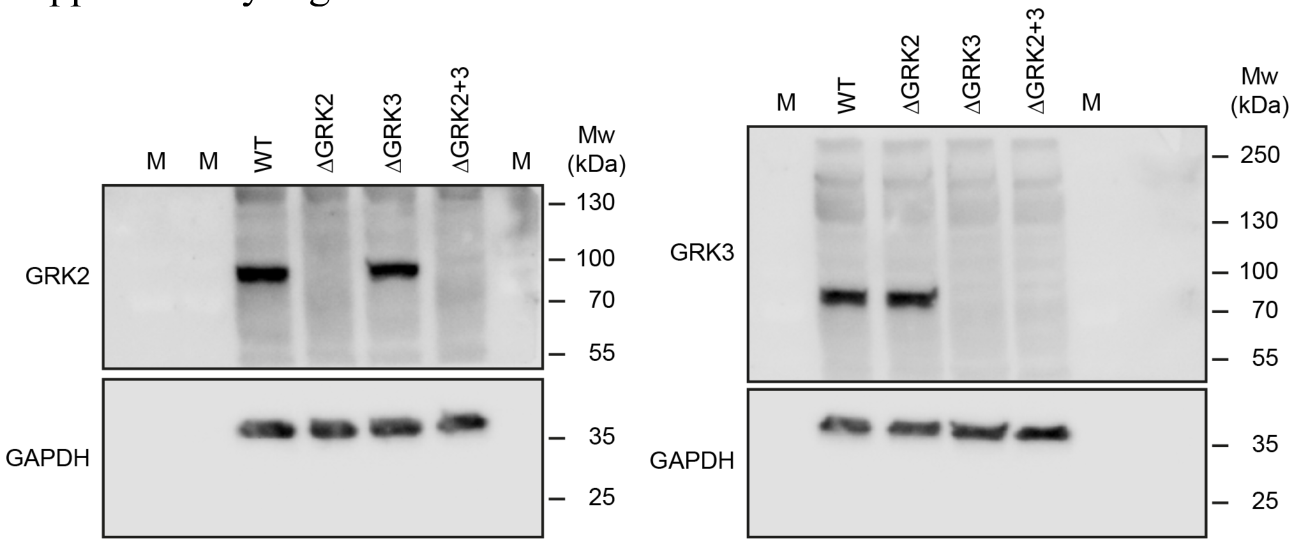


**Supplementary Figure S1.** Full-length blots corresponding to the cropped blots in Fig. 1a. M, protein marker.

### Supplementary Figure S2

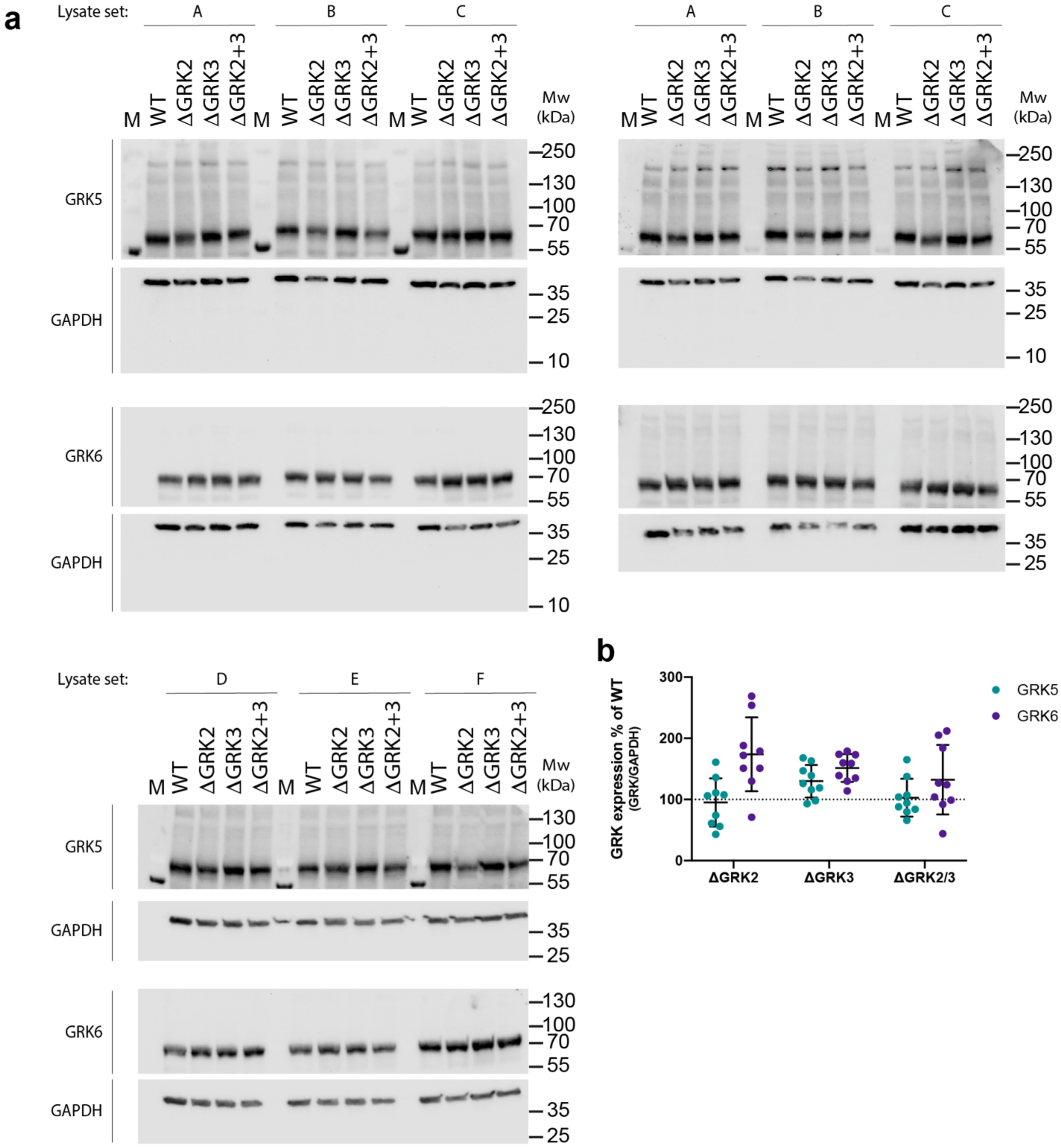


**Supplementary Figure S2.** Protein levels of GRK5 and GRK6 in parental and genome-edited HEK293A cells. (**a**) Western blot evaluation of GRK5 or GRK6 expression in parental HEK293A cells and ΔGRK2, ΔGRK3 or ΔGRK2/3 cell lines. GAPDH levels serve as loading control. Six sets of cell lysate preparations (A-E) were analyzed in three western blot experiments (all blots are shown). (**b**) Quantification of GRK5 and GRK6 levels after correction to the GAPDH control and normalization to the WT sample. Data points represent quantification of individual lanes depicted in (a) and the line and error bars represent the mean ± SD.

### Supplementary Figure S3

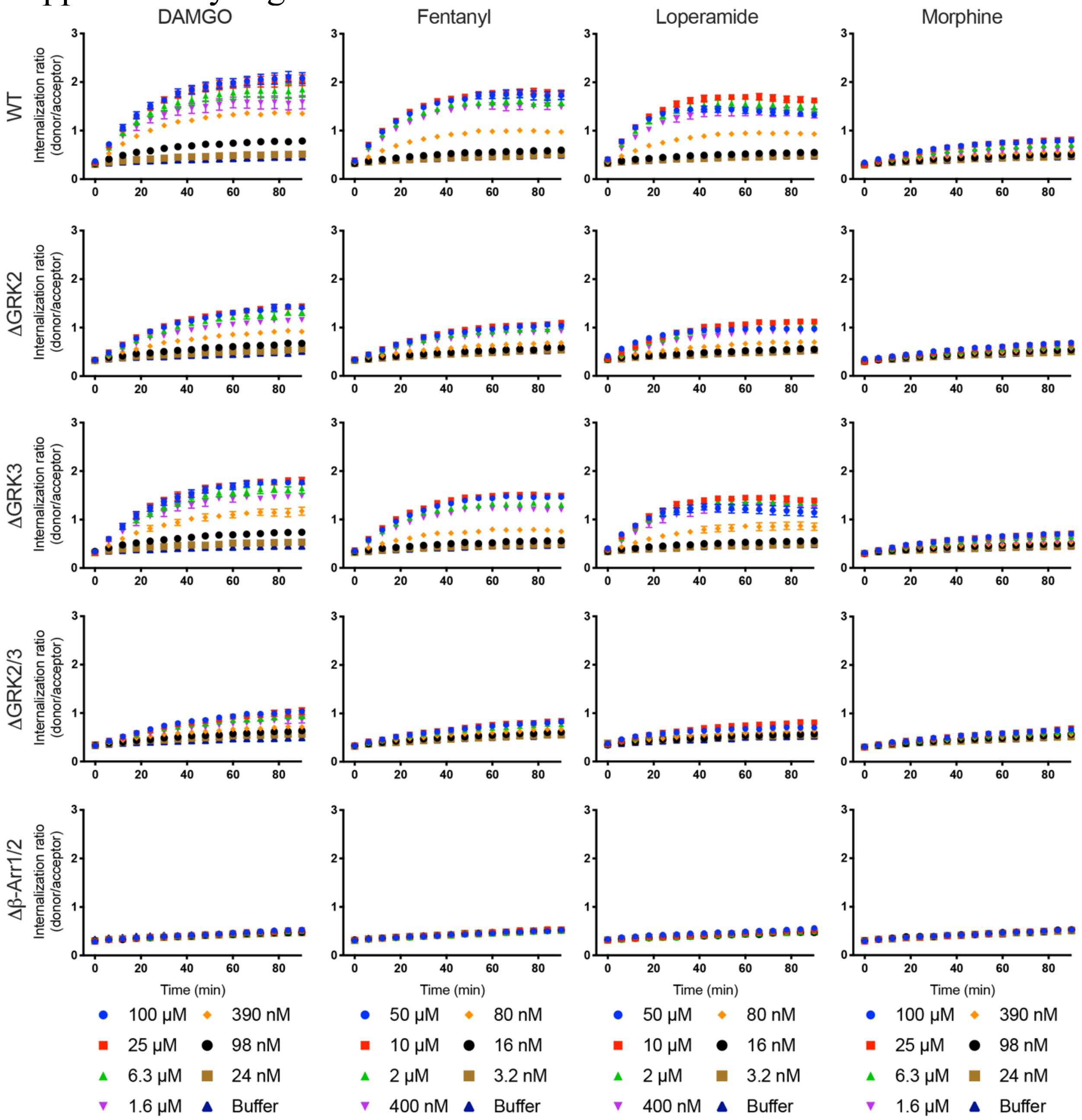


**Supplementary Figure S3.** Real-time internalization curves of µ-OR in parental or genome-edited cell lines in the absence of agonist (buffer) or stimulated by a range of concentrations of DAMGO, loperamide, fentanyl or morphine. Data represent the mean ± SEM of the internalization ratio (donor signal/acceptor signal) from 3-5 independent experiments carried out in duplicate. Error bars not shown lie within the dimension of the symbol.

### Supplementary Figure S4

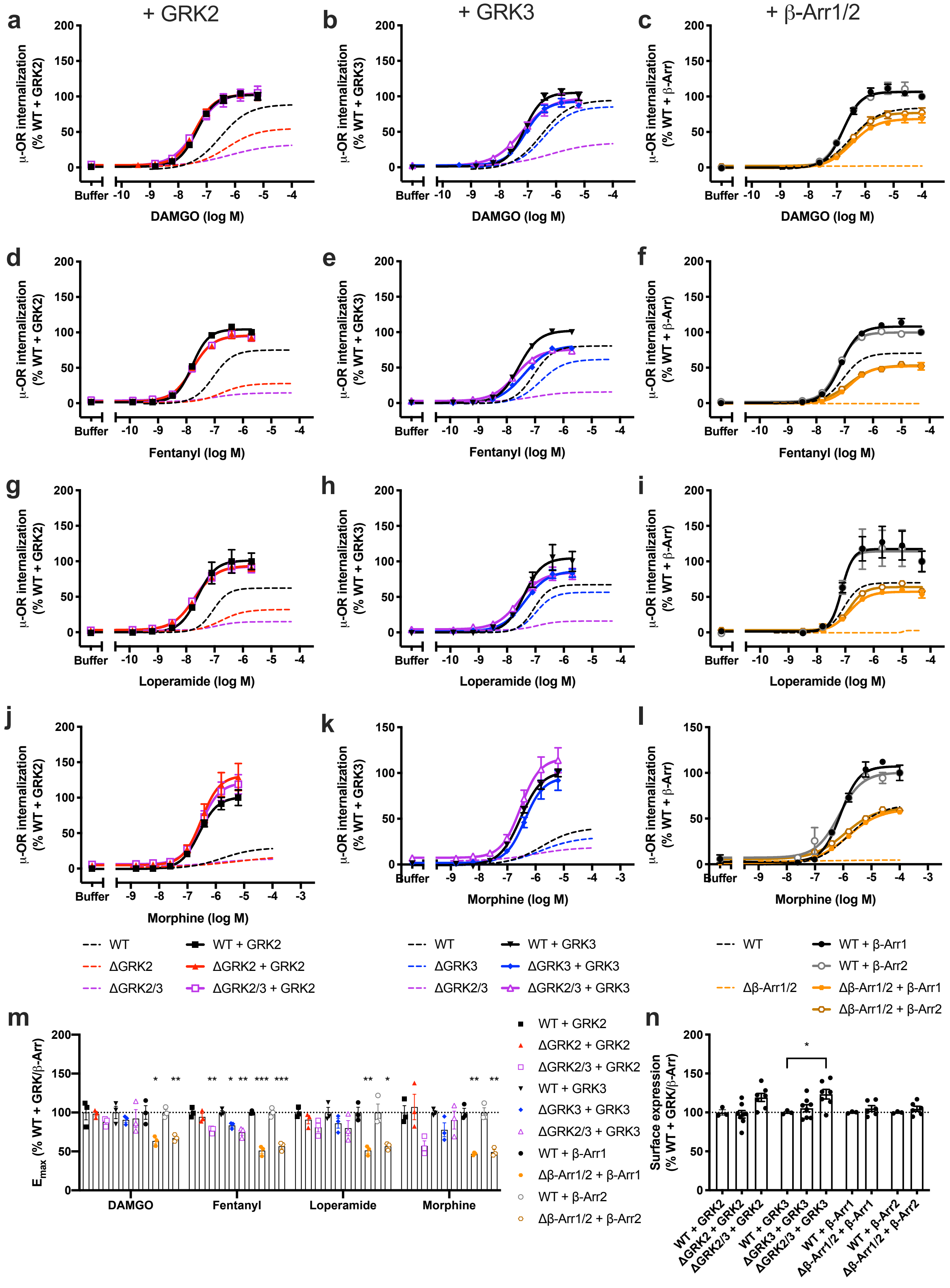

**Supplementary Figure S4.** μ-OR internalization in genome-edited cell lines with overexpression of the deleted proteins. Concentration-response curves from stimulation with (**a-c**) DAMGO, (**d-f**) fentanyl, (**g-i**) loperamide, or (**j-l**) morphine. Internalization without overexpression is shown for comparison (dashed lines). Data represent the mean ± SEM of the area under the curve of 90 min real-time internalization experiments after subtraction of the buffer response in parental (WT) cells and normalization to the maximum response in the parental cells with overexpression of the same protein from 3-8 independent experiments carried out in duplicate. Error bars not shown lie within the dimension of the symbol. (**m**) $E_{max}$ of agonist responses in parental and genome-edited cells with overexpression of GRK2/3 or β-arrestin1/2. (**n**) μ-OR cell surface expression in parental and genome-edited cell lines with overexpression of the deleted proteins. (**m-n**) Data represent the mean of individual experiments (circles) as well as mean ± SEM (columns) after normalization to expression in the parental cells with overexpression of the same protein from 3 (**m**) or 3-8 (**n**) independent experiments with 2 (**m**) or 32 (**n**) replicates per experiment. Values in genome-edited cells with overexpression was compared to the parental cells with overexpression of the same protein by one-way ANOVA with Dunnett's multiple comparisons test (GRK2/3 overexpression) or unpaired t-test (β-arrestin1/2 overexpression). *$P = 0.01$-$0.05$, **$P = 0.001$-$0.01$, ***$P < 0.001$.

### Supplementary Figure S5

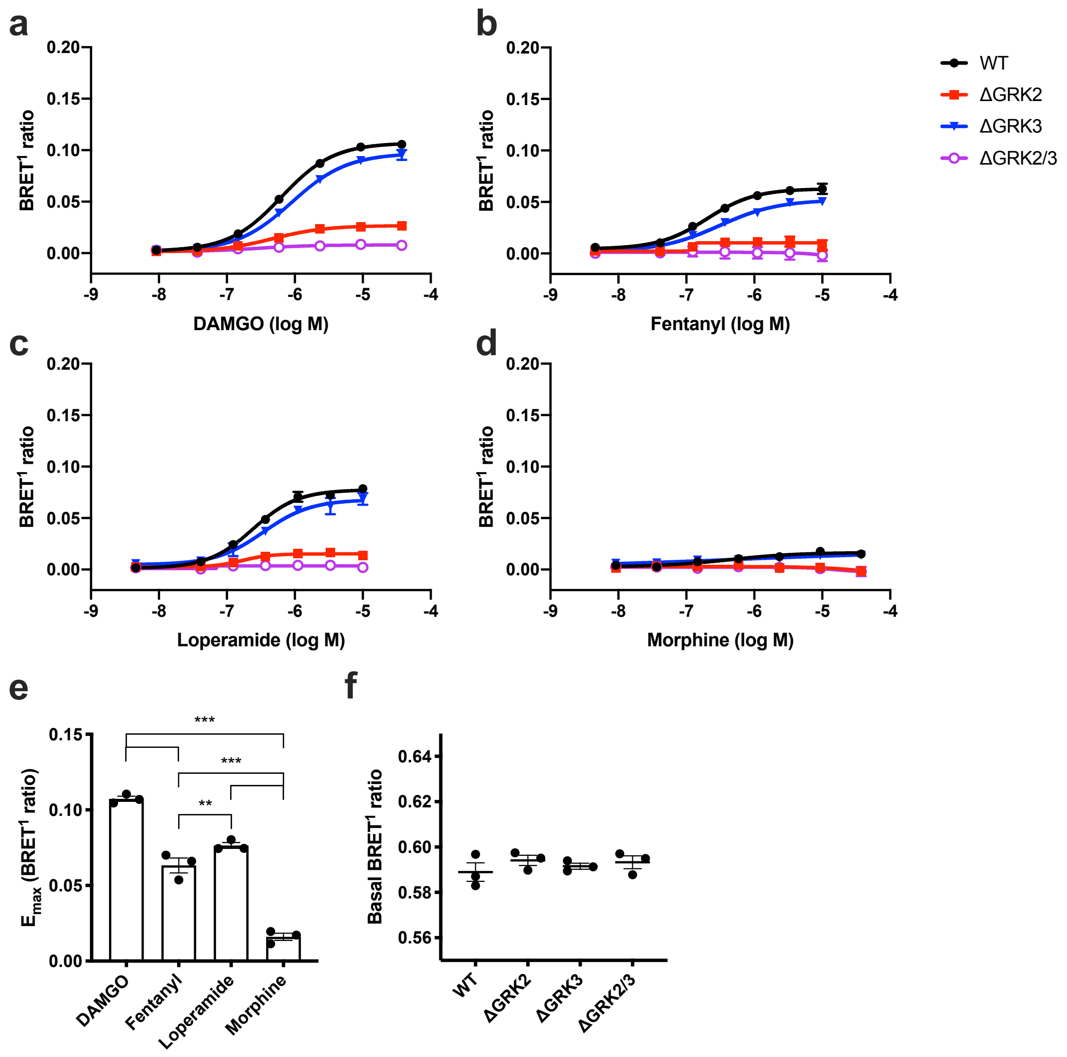


**Supplementary Figure S5.** β-arrestin2 recruitment of μ-OR in genetically modified cell lines. Recruitment of β-arrestin2-RlucII to μ-OR-EYFP after 6 min stimulation of μ-OR by a range of concentrations of (**a**) DAMGO, (**b**) fentanyl, (**c**) loperamide, or (**d**) morphine in parental HEK293A (WT) cells or HEK293A cell lines with deletion of GRK2 (ΔGRK2), GRK3 (ΔGRK3), or GRK2 and -3 (ΔGRK2/3). Data represent the mean ± SEM of the $BRET^1$ ratio after subtraction of the buffer response in parental cells from 3-4 independent experiments carried out in duplicate. Error bars not shown lie within the dimension of the symbol. (**e**) $E_{max}$ of agonist responses in parental cells. Data represent the mean of individual experiments (circles) and the mean ± SEM (columns) from 3 individual experiments. Statistical comparison by one-way repeated measures ANOVA with Dunnett's multiple comparisons test. $**P = 0.001\text{-}0.01$, $***P < 0.001$. (**f**) Basal β-arrestin2 recruitment determined as the buffer response. Data represent the mean of individual experiments (circles) and the mean ± SEM (lines) from 3 independent experiments with 8 replicates per experiment. Basal recruitment in knockout cells was compared with parental cells by one-way repeated measures ANOVA with Dunnett's multiple comparisons test and no significant differences were found ($P > 0.05$).

### Supplementary Figure S6

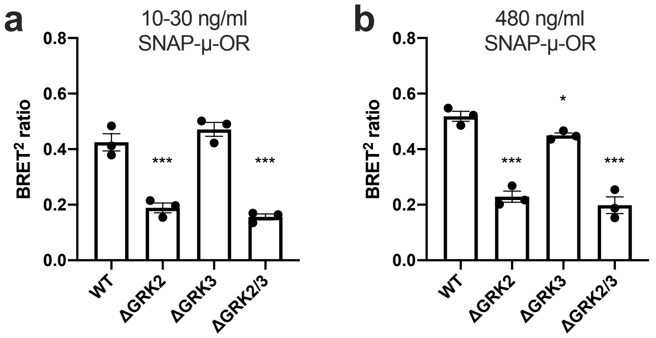


**Supplementary Figure S6.** Increasing the μ-OR DNA amount does not change the β-arrestin2 recruitment pattern in the ebBRET assay with 6 min stimulation by 37.5 μM DAMGO. Parental and GRK knockout cells were transfected with 10-30 ng/ml SNAP-μ-OR (**a**) or 480 ng/ml SNAP-μ-OR (corresponding to the μ-OR DNA amount in the μ-OR-EYFP β-arrestin2 recruitment assay) (**b**). Data represent the mean of individual experiments (circles) and mean ± SEM (columns) from 3 independent experiments performed in duplicate. BRET ratios in knockout cells were compared with parental cells by repeated measures one-way ANOVA with Dunnett's multiple comparisons test. *$P$ = 0.01-0.05, ***$P$ < 0.001.

### Supplementary Figure S7

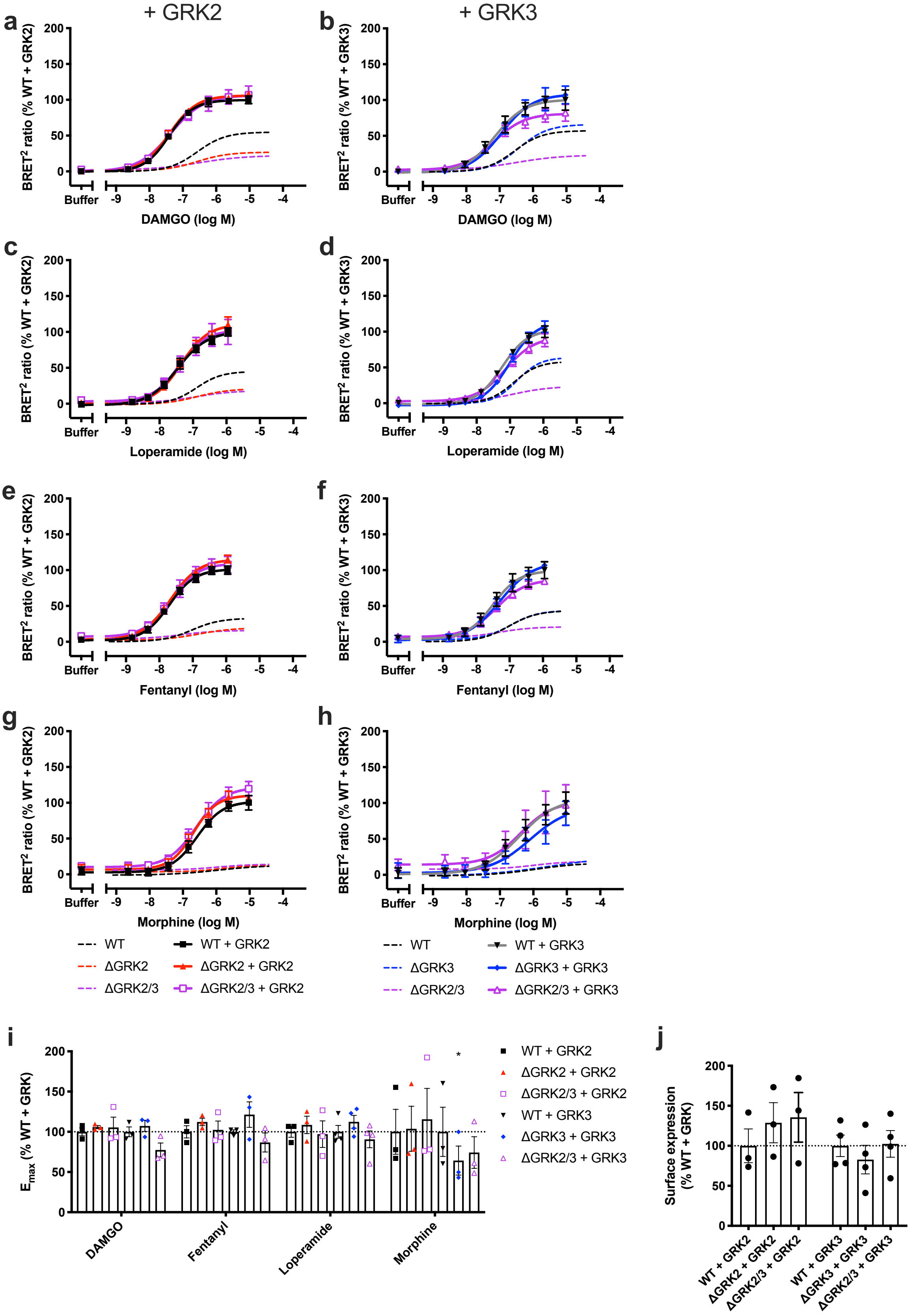

**Supplementary Figure S7**. β-arrestin2 recruitment by μ-OR in genome-edited cell lines with overexpression of the deleted GRKs. β-arrestin2 recruitment was measured by ebBRET after 6 min stimulation of μ-OR with (**a-b**) DAMGO, (**c-d**) fentanyl, (**e-f**) loperamide, or (**g-h**) morphine. β-arrestin2 recruitment without GRK overexpression is shown for comparison (dashed lines). Data represent the mean ± SEM after subtraction of the buffer response in parental (WT) cells and normalization to the maximum response in the parental cells with overexpression of the same protein from 3-4 independent experiments carried out in duplicate. Error bars not shown lie within the dimension of the symbol. (**i**) $E_{max}$ of agonist responses in parental and genome-edited cells with overexpression of GRK2 or -3. (**j**) μ-OR cell surface expression determined with ELISA. (**i-j**) Data represent the mean of individual experiments (circles) and mean ± SEM (columns) after normalization to expression in the parental cells from 3-4 independent experiments with 2 (**i**) or 3 (**j**) replicates per experiment. Values in genome-edited cells with overexpression was compared to the parental cells with overexpression of the same protein by one-way ANOVA with Dunnett's multiple comparisons test. *$P$ = 0.01-0.05.

#### Supplementary Figure S8

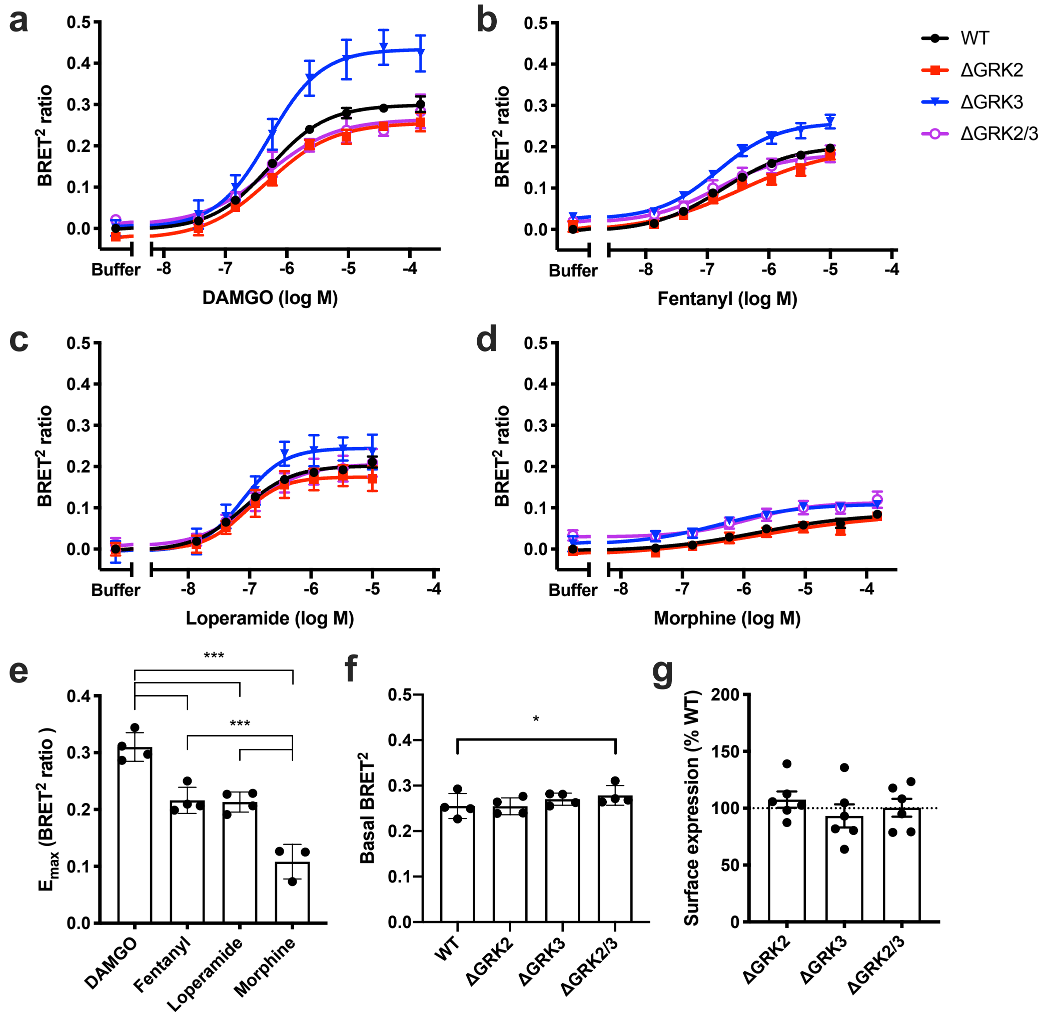


**Supplementary Figure S8.** β-arrestin2 recruitment of μ-OR in genetically modified cell lines. Recruitment of β-arrestin2-RlucII to the cell membrane monitored by ebBRET after 60 min stimulation of μ-OR by a range of concentrations of (**a**) DAMGO, (**b**) fentanyl, (**c**) loperamide, or (**d**) morphine in parental HEK293A (WT) cells or HEK293A cell lines with deletion of GRK2 (ΔGRK2), GRK3 (ΔGRK3), or GRK2 and -3 (ΔGRK2/3). Data represent the mean ± SEM of the $BRET^2$ ratio after subtraction of the buffer response in parental cells from 3-4 independent experiments carried out in duplicate. Error bars not shown lie within the dimension of the symbol. (**e**) $E_{max}$ of agonist responses in parental cells. Statistical comparison by one-way ANOVA with Dunnett's multiple comparisons test. ***$P < 0.001$. (**f**) Basal β-arrestin2 recruitment determined as the buffer response. (**g**) μ-OR surface expression measured by ELISA and normalized to expression in parental cells. (**e-**

**g**) Data represent the mean of individual experiments (circles) and the mean ± SEM (columns) from 3-4 (**e**) 4 (**f**) or 6 (**g**) independent experiments with 2 (**e**), 4-8 (**f**) or 3 (**g**) replicates per experiment. (**f-g**) Values in knockout cells were compared with parental cells by one-way ANOVA with Dunnett's multiple comparisons test. *$P$ = 0.01-0.05.

### Supplementary Figure S9

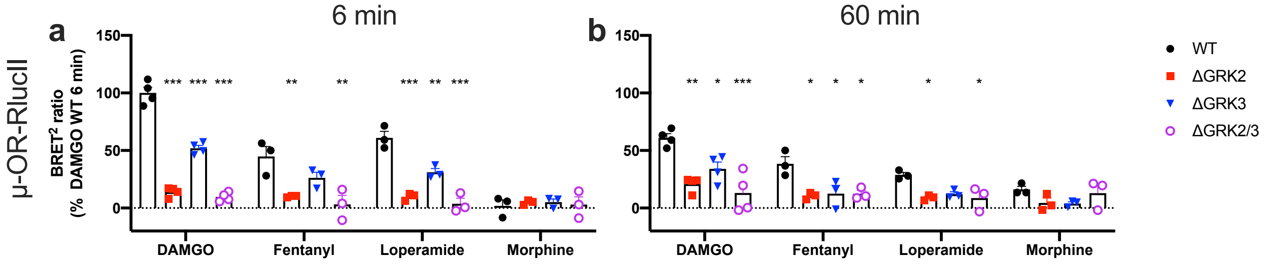


**Supplementary Figure S9.** Time dependence of β-arrestin1 recruitment of μ-OR in ΔGRK cell lines. Recruitment of GFP$^2$-β-arrestin2 to μ-OR-RlucII after 6 min (**a**) or 60 min (**b**) stimulation by saturating concentrations of DAMGO (37.5 μM), fentanyl (10 μM), loperamide (10 μM) or morphine (37.5 μM). Data represent the mean of individual experiments (circles) as well as the mean ± SEM (columns) after buffer subtraction and normalization to the maximum response in parental (WT) cells stimulated with DAMGO for 6 min from 3-4 independent experiments performed in duplicate. BRET ratios in knockout cells were compared with parental cells by repeated measures one-way ANOVA with Dunnett's multiple comparisons test. **P* = 0.01-0.05, ***P* = 0.001-0.01, ****P* < 0.001.

### Supplementary Figure S10

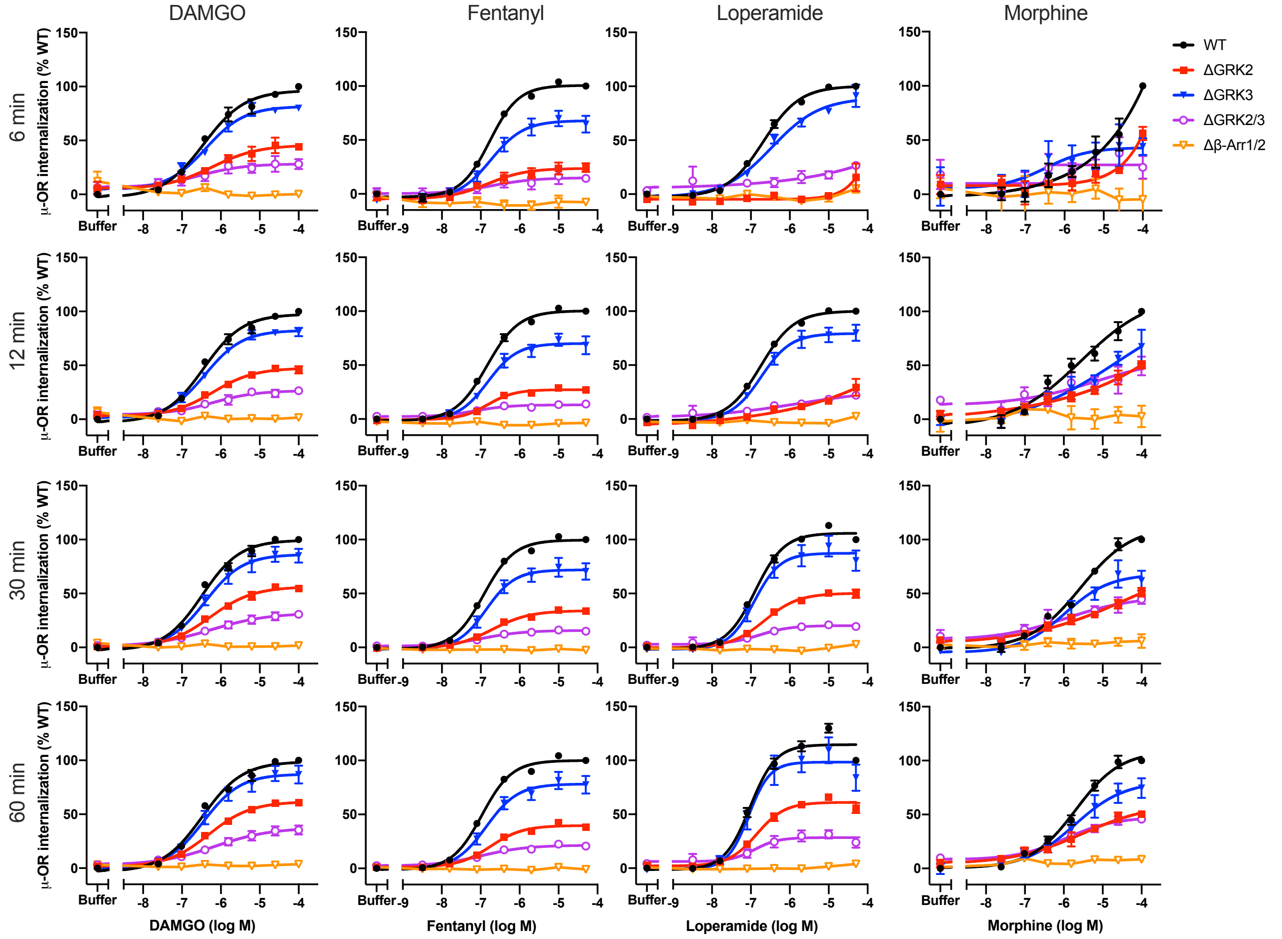


**Supplementary Figure S10.** μ-OR internalization concentration-response curves in genome-edited cell lines with different stimulation times. Concentration-response curves from stimulation of μ-OR internalization with a range of concentrations of DAMGO, fentanyl, loperamide and morphine plotted using the response after 6, 12, 30 or 60 min stimulation. Data represent the mean ± SEM after subtraction of the buffer response in parental (WT) cells and normalization to the maximum response in parental cells from 3-5 independent experiments performed in duplicate. Error bars not shown lie within the dimension of the symbol.

**Supplementary Table S1.** $E_{max}$ (% of parental HEK293A (WT) cells) and $pEC_{50}$ values for recruitment of β-arrestin2-RlucII to μ-OR-EYFP measured by BRET[1] after 6 min of μ-OR stimulation. The values were obtained from fitting concentration-response curves to a four-parameter model of agonism. The responses were too low in ΔGRK2 cells with fentanyl or morphine as agonist and in ΔGRK2/3 cells to fit the data (n.d., not determined). $E_{max}$ and $pEC_{50}$ values are the mean of 3 independent experiments. $E_{max}$ and $pEC_{50}$ in knockout cells were compared to the parental cells before normalization by either one-way repeated measures ANOVA with Dunnett's multiple comparisons test (DAMGO and loperamide) or paired t-test (fentanyl and morphine). $^{*}P = 0.01\text{-}0.05$, $^{***}P < 0.001$.

| | WT | | | | ΔGRK2 | | | | ΔGRK3 | | | | ΔGRK2/3 | | | |
|---|---|---|---|---|---|---|---|---|---|---|---|---|---|---|---|---|
| | $E_{max}$ | SEM | $pEC_{50}$ | SEM | $E_{max}$ | SEM | $pEC_{50}$ | SEM | $E_{max}$ | SEM | $pEC_{50}$ | SEM | $E_{max}$ | SEM | $pEC_{50}$ | SEM |
| DAMGO | 100 | 2 | 6.21 | 0.04 | $25^{***}$ | 3 | $6.36^{*}$ | 0.03 | 92 | 7 | $6.04^{*}$ | 0.03 | n.d. | | n.d. | |
| Fentanyl | 100 | 11 | 6.75 | 0.03 | n.d. | | n.d. | | 82 | 8 | 6.53 | 0.13 | n.d. | | n.d. | |
| Loperamide | 100 | 4 | 6.61 | 0.02 | $19^{***}$ | 6 | 6.81 | 0.09 | 91 | 17 | 6.47 | 0.01 | n.d. | | n.d. | |
| Morphine | 100 | 21 | 6.65 | 0.25 | n.d. | | n.d. | | 79 | 17 | 7.44 | 0.39 | n.d. | | n.d. | |

**Supplementary Table S2.** $E_{max}$ (% of parental HEK293A (WT) cells) and $pEC_{50}$ values for β-arrestin2-RlucII recruitment to the plasma membrane measured by ebBRET after 60 min of μ-OR stimulation. The values were obtained from fitting concentration-response curves to a four-parameter model of agonism. $E_{max}$ and $pEC_{50}$ values are the mean of 3-4 independent experiments. $E_{max}$ and $pEC_{50}$ in knockout cells were compared to the parental cells by one-way ANOVA with Dunnett's multiple comparisons test. $^{*}P = 0.01\text{-}0.05$.

| | WT | | | | ΔGRK2 | | | | ΔGRK3 | | | | ΔGRK2/3 | | | |
|---|---|---|---|---|---|---|---|---|---|---|---|---|---|---|---|---|
| | $E_{max}$ | SEM | $pEC_{50}$ | SEM | $E_{max}$ | SEM | $pEC_{50}$ | SEM | $E_{max}$ | SEM | $pEC_{50}$ | SEM | $E_{max}$ | SEM | $pEC_{50}$ | SEM |
| DAMGO | 100 | 6 | 6.28 | 0.06 | 92 | 5 | 6.30 | 0.02 | 140* | 6 | 6.29 | 0.02 | 84 | 13 | 6.30 | 0.06 |
| Fentanyl | 100 | 8 | 6.72 | 0.13 | 97 | 20 | 6.52 | 0.19 | 108 | 9 | 6.84 | 0.09 | 78 | 9 | 6.83 | 0.08 |
| Loperamide | 100 | 6 | 7.11 | 0.02 | 84 | 8 | 7.07 | 0.09 | 117 | 5 | 7.11 | 0.04 | 94 | 11 | 6.99 | 0.11 |
| Morphine | 100 | 23 | 5.70 | 0.29 | 89 | 25 | 5.68 | 0.65 | 123 | 27 | 5.93 | 0.49 | 88 | 29 | 5.94 | 0.24 |
